# Afanc: a Metagenomics Tool for Variant Level Disambiguation of NGS Datasets

**DOI:** 10.1101/2023.10.05.560444

**Authors:** Arthur V. Morris, Anna Price, Tom Connor

## Abstract

Genomics is amongst the most powerful tools available for mounting a clinical response to infectious disease. The accurate and precise taxonomic evaluation of pathogens is essential when building a picture of pathogenicity, virulence, transmission, and drug resistance. Carrying out such profiling in a high throughput manner necessitates the development of reliable bioinformatic tools. Here we present Afanc, a novel metagenomic profiler which is sensitive down to species and strain level taxa, and capable of elucidating the complex pathogen profile of compound datasets. We compared Afanc against currently available cutting edge profilers using 3 datasets: single species read sets simulated from the full Mycobacteriaceae taxonomic landscape; compound read sets containing multiple Mycobacteriaceae species and variants; and real data covering the majority of the *M. tuberculosis* lineage taxonomic space. Afanc outperformed all profilers, both generic and Mycobacteriaceae specific, across all tested fields. As a species agnostic profiler, we predict that Afanc will be of great utility when carrying out highly specific and sensitive pathogen profiling of clinical datasets. Such analyses are essential in advising both the clinical response to an individual disease case, and in forming the foundation of epidemiological surveys.

## 1. Background

The need for reliable and robustly tested genomic and metagenomic profilers cannot be understated. Such profilers form the backbone of speciation functionality within many bioinformatic pipelines utilised by medical laboratories and public health agencies. Kraken1 & 2 (Wood and Salzberg, 2014; Wood et al., 2019) and species-level sequence abundance estimation algorithms building upon the Kraken framework, such as Bracken (Lu et al., 2017) and KrakenUniq (Breitwieser et al., 2018) have proven to be effective in the disambiguation of genomic and metagenomic datasets. However, these tools may not be sufficient, on their own, for use in settings where treatment decisions will be dictated by the results they produce. In recent years, Mycobacteriaceae specific profilers such as Mykrobe (Hunt et al., 2019) and TB-Profiler (Napier et al., 2020) have led the way in performing species and lineage level identification of Mycobacteriaceae, aiming to provide more robust speciation, suitable for both research and clinical/public health laboratory use. Sensitive and reliable taxonomic designation is an essential task in the domain of public health, both informing the clinical response to an individual infection, and forming the foundation of epidemiological surveys.

Generally, two approaches are taken to metagenomic species/lineage identification.

1) Genomic distance based: the tool assigns a taxonomic ID based on genomic distance from a set of genome assemblies within a database.
2) Variant profile based: the tool screens against a set of lineage defining canonical mutations.

Genomic distance based tools, such as Kraken1 & 2, are optimised for identifying species level taxa within complex metagenomic NGS read sets. NGS reads represent only a very small fragment of the overall genome. Their relatively small size, and consequent limited sequence specificity, results in a large number of false positive taxonomic assignments. This results in the reports produced by these tools often being very complex, with a large and diverse range of reported taxa. Simplification of such reports is of primary concern to derivative tools. Bracken achieves this by using Bayes rule to re-estimate the distribution of reads across taxa of a given rank from their children taxa. This gives a more precise and robust measure of taxonomic abundance within a read set. KrakenUniq utilises *k*-mer cardinality estimation to reduce false-negative (FN) and false-positive (FP) taxonomic assignment of reads exhibited in Kraken reports, giving a clearer picture of the true genetic space covered by the DNA in the sample. Both of these approaches use the *k*-mer content of the sequence spaces occupied by each taxonomic rank to optimise read binning, and consequently, improve reporting precision. This approach relies on there being sufficient distance between two discrete taxonomic sequence spaces to identify FP and FN assignments. Where the genomic distance between two taxonomic ranks is very low (such as in the case of TB lineages), the sequence space they occupy may be too small to allow for reliable reclassification. Consequently, such approaches may only be reliable down to species and subspecies level, and alternative approaches must be utilised to perform higher resolution taxonomic identification. Profiling tools such as Mykrobe and TB-Profiler solve this problem by identifying variant defining mutations within a given species sequence. Mykrobe utilises probes, short DNA sequences of length 2*k* − 1, within which a lineage or species defining mutation is embedded, such that a probe represents a SNP with sequences of *k* bases flanking it. TB-Profiler identifies key SNPs within a read set and screens them against a profile of known lineage defining SNPs. Both of these approaches are highly sensitive, but require taxonomic ranks to be defined by discriminatory sets of mutations. Construction of these sets may be prohibitively time consuming for large numbers of taxa, where the mutation space is very large.

There are currently just under 200 species of Mycobacteriaceae described. For the purpose of clinical practicality, these are generally divided into two categories: Tuberculous (TB-complex) and Non-Tuberculous Mycobacteriacea (NTM). In 2018, based on core genome phylogenetic analysis, the 188 species comprising the genus Mycobacterium at that time were divided into 5 distinct genera: Mycobacterium, Mycobacteroides, Mycolicibacillus, Mycolicibacter, and Mycolicibacterium (Gupta et al., 2018). *Mycobacterium tuberculosis* itself is divided into lineages, which are defined by the presence of key single nucleotide polymorphisms (SNPs). Lineage specific genomic diversity is known to have influence on virulence, transmissibility, drug resistance, and host response (Napier et al., 2020). Reliable identification of lineage is therefore a fundamental factor when determining the pathogenic profile of *Mycobacterium tuberculosis* cases.

Bacteria belonging to Mycobacteriaceae are of enormous clinical importance, with over 2000 samples passing through the Wales Centre for Mycobacteria (WCM) alone every year. Prevalence of TB globally is estimated by the WHO to be 10.6 million active cases a year, resulting in 1.6 million deaths (WHO TB report 2022), with an estimated 1.7 billion latent infections (Houben and Dodd, 2016). Furthermore, global incidence and deaths from NTM diseases have been steadily rising. Despite NTMs previously being thought of as exclusively opportunistic pathogens of the immunocompromised, infections in immunocompetent individuals are being reported at an increasing rate. Such infections are often highly drug resistant, and present with complex pathology which resists treatment (Ratnatunga et al., 2020).

Here, we present Afanc, a taxonomic profiler capable of both species and lineage level identification. We solve the issues detailed above by carrying out species and subspecies level profiling using a novel Kraken2 report disambiguation algorithm, and lineage level profiling using a variant profiling approach. We demonstrate that this hybrid approach results in a “best of both worlds” outcome, whereby both researchers and clinical/public health labs can benefit from the speed and reliability of species level identification by genomic distance, and the sensitivity of lineage level identification using variant profiles.

## 2. Implementation

Afanc consists of 3 discrete sub-tools: `get_dataset`, `autodatabase`, and `screen`. These modules function both as standalone tools, and are designed to integrate to form a single workflow.

### 2.1 get_dataset

This module downloads a dataset of genome assemblies from Genbank belonging to species defined within a text file. Genome assemblies are downloaded and deposited in a directory structure consisting of a parent directory containing subdirectories for each species. Subspecies and variants specified within the text file are deposited in their own subdirectory within their parent species directory. This directory structure can be used as input for the `autodatabase` module.

For example, given the text file shown in Figure 1, and the user defined number of assemblies for each ID to download as 3, the structure of the output directory will be that seen in Figure 2.

**Figure 1.**
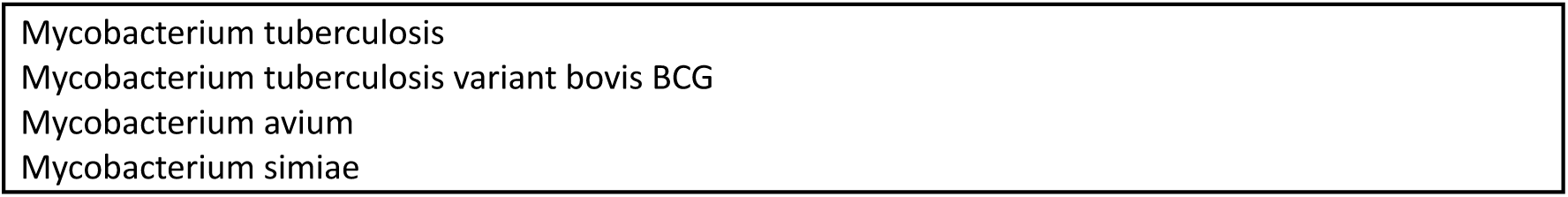
The structure of a text file used as input for the Afanc get_dataset module.

**Figure 2.**
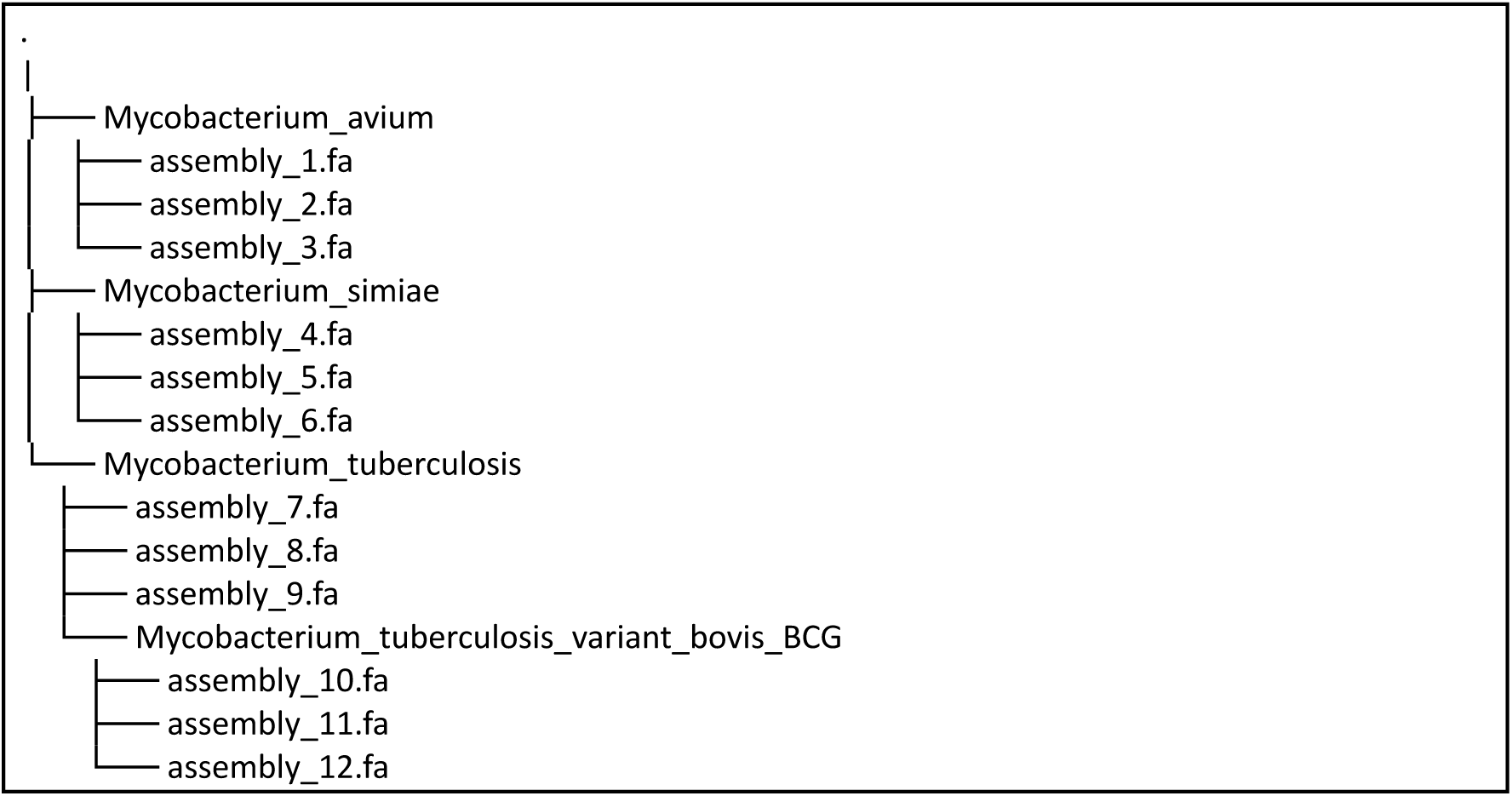
The output directory structure produced by running the Afanc get_dataset module using the text file outlined in fig. 1.

### 2.2 autodatabase

The `autodatabase` module automates the process of quality control and database construction. Naive database creation from sequences downloaded from NCBI often results in the inclusion of poor quality sequences. This module is designed to deal with this problem by performing quality control of input sequences, whilst also allowing users to update databases as new sequences are made available. The `autodatabase` module takes genome assemblies contained within a directory structure of the form generated by the `get_dataset` module (see Figure 2). This directory structure must contain directories for each species level taxon, where subdirectories within each species directory pertain to subspecies and variant (here referring to taxa lower than subspecies) level taxa, or any other taxonomic rank lower than species.

There are 6 primary stages to the workflow of this module, which can be seen in Figure 3. First, the NCBI taxonomy database from the date specified by the user is downloaded (step 1). By default, this will be the database from 2022-05-01. The NCBI taxonomy database is effectively a tree, where each taxonomic rank refers to a node within this tree and is assigned a numeric taxonomy ID. If a taxon named within the input directory structure is not assigned to a node within the NCBI database, Afanc will attempt to assign it a taxonomy ID and create a simulated node. This process will fail if the named taxon cannot be assigned to a known genus. A Mash matrix is constructed from genome assemblies belonging to each specified taxon (step 2) (Ondov et al., 2016, Napier et al., 2020). This Mash matrix is used to select the highest quality assemblies for database construction, by filtering out assemblies which lie outside a given range (by default, this is 0.1) of the mode of the average mash distance for all assemblies within that taxon (step 3). This is to ensure that low quality assemblies are removed, to prevent erroneous screening results. The Mash matrix is defined as follows

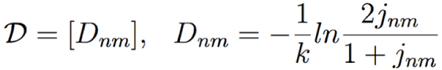

**Figure 3.**
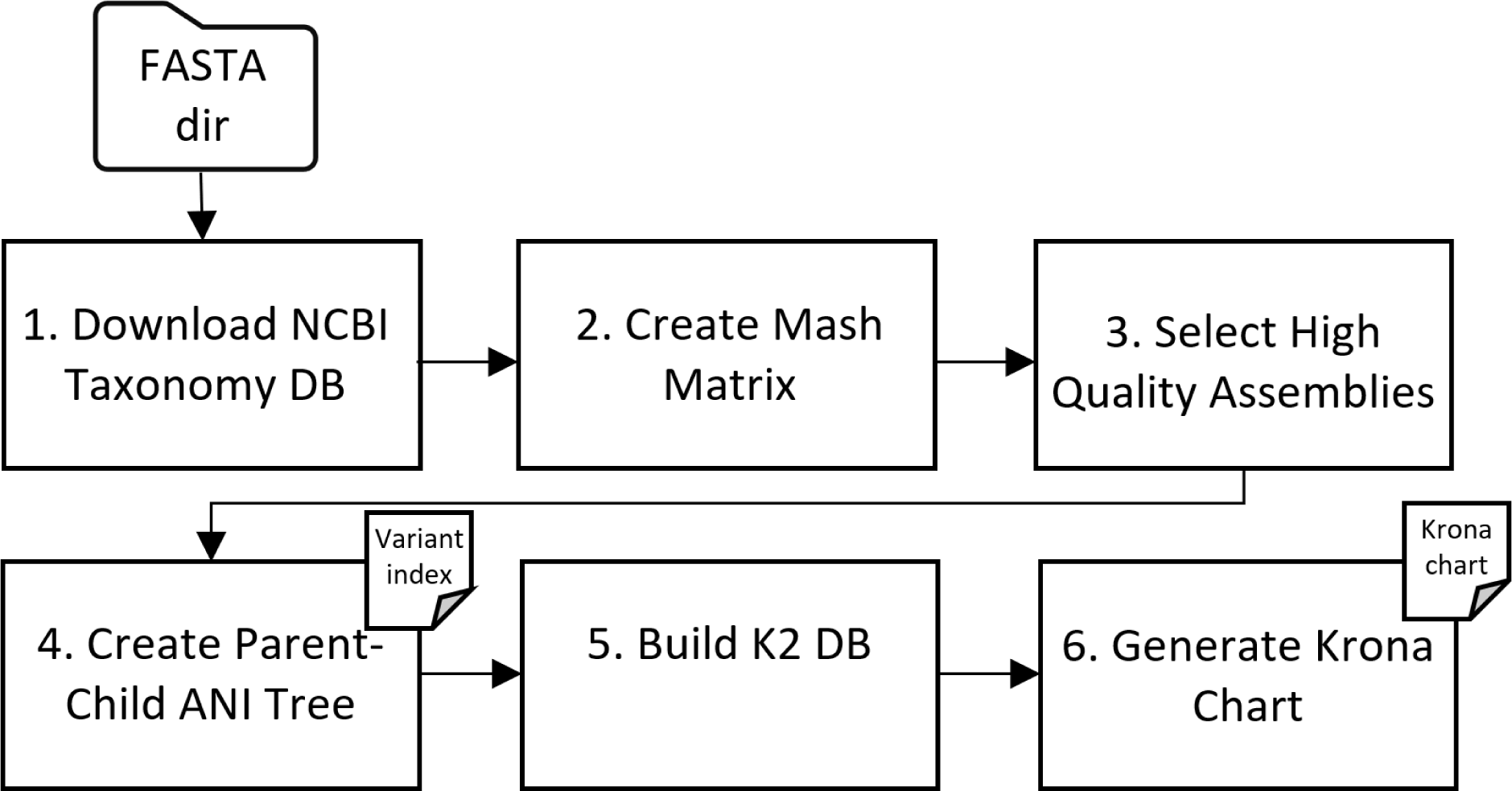
The autodatabase module workflow.

where *D_nm_* is the pairwise distance between samples *n* and *m*, *k* is the size of the mash sketch and *j* is the Jaccard estimate.

Parent-child taxon average nucleotide identity (ANI) distances are calculated from the set of high-quality assemblies (step 4). This is achieved by iterating through each taxon and calculating both the intrataxon ANI, and ANI of the assemblies within the taxon and its parent taxon. These results are stored in a JSON file and used to calculate the elastic threshold for each taxon (see Section 2.3.1.1). A Kraken 2 (K2) database is then constructed using these high-quality assemblies (step 5). Finally, a Krona chart is generated for easy visualisation of the database (step 6).

The output directory from this module constitutes the database used for running the Afanc `screen` module. It consists of a directory containing 5 JSON files, and 6 subdirectories (see figure 4). The K2 database is contained within the krakenBuild_autoDatabase_kraken2Build subdirectory. The Krona chart for visualisation of the assemblies used to construct the K2 database can be found within the krakenBuild_autoDatabase_krona subdirectory.

**Figure 4.**
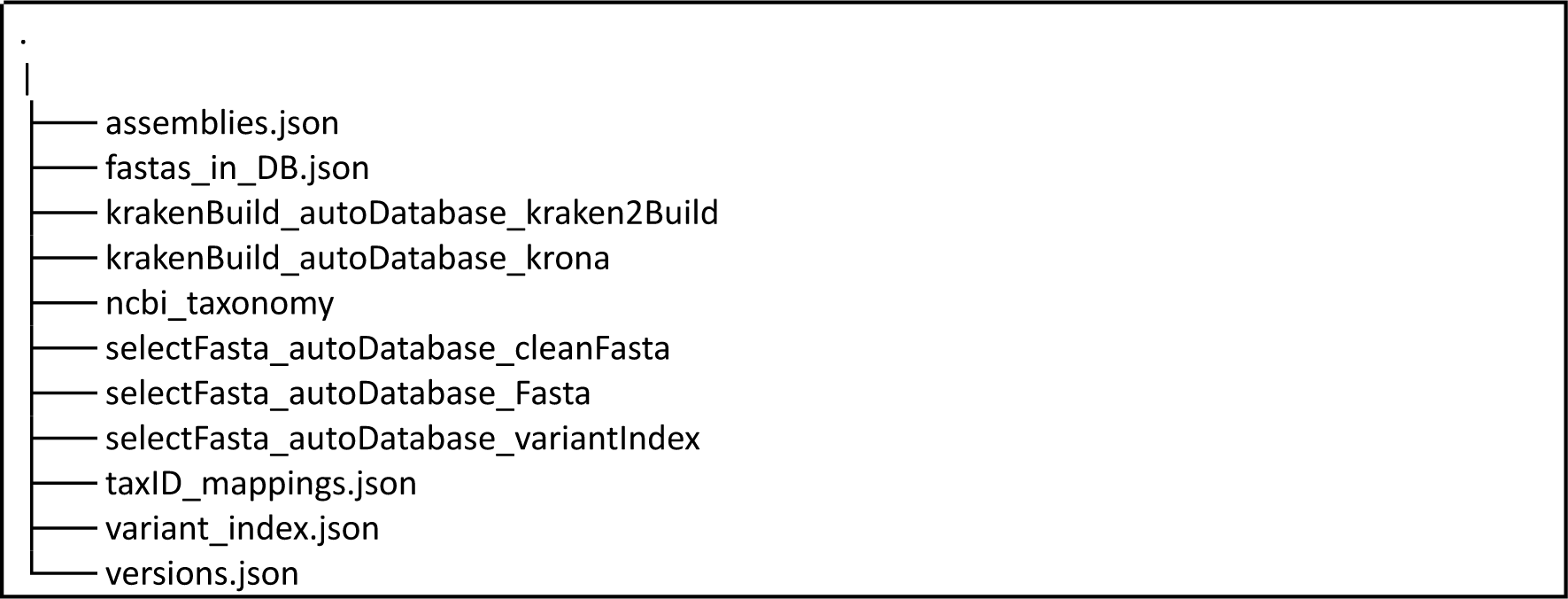
The output directory structure produced by running the Afanc autodatabase module using the directory structure seen in fig 3.

**Figure 5.**
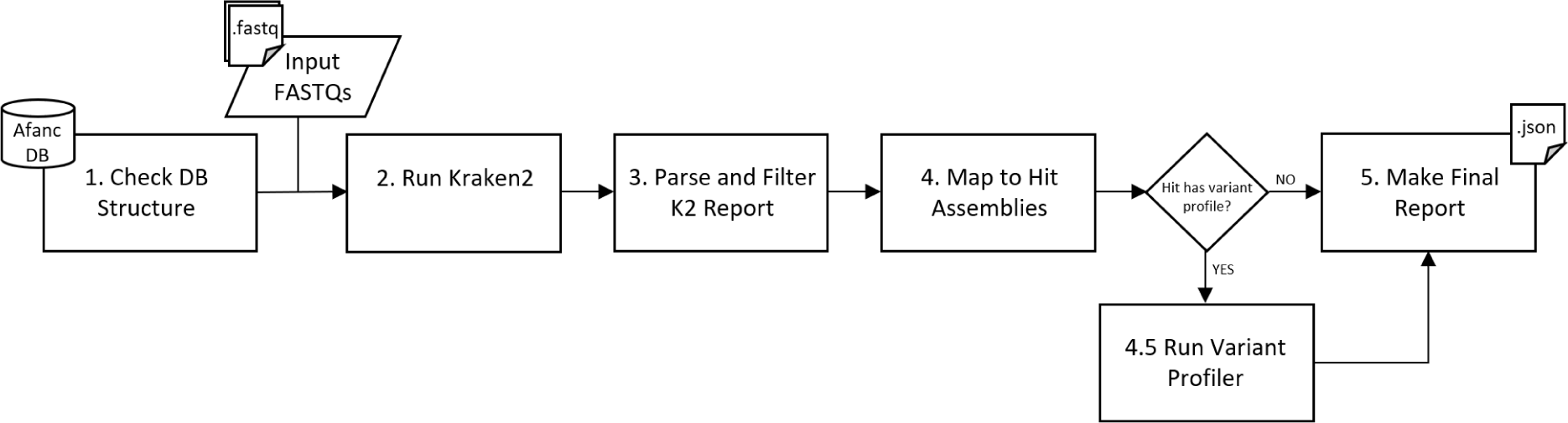
The screen module workflow.

### 2.3 screen

The `screen` module performs a metagenomic survey of a given read set. It takes a database created by running Afanc `autodatabase` (see figure 4), and a set of reads in FASTQ format. A JSON format report is produced, detailing the metagenomic profile of the input read set.

There are five principal stages to the workflow of this module. First, the input database is checked to ensure it is not malformed (step 1). If this check is passed, Kraken 2 is used to produce a general metagenomic report of the input read set, using the krakenBuild_autoDatabase_kraken2Build input database subdirectory as the K2 database (step 2). The output K2 report is then parsed using a novel algorithm to identify the most likely species and variants within the input read set (step 3). A full explanation for the algorithmic approach to solving this problem can be found in Section 2.3.1. The input read set is then subjected to a competitive mapping protocol, where reads are mapped to the repertoire of genome assemblies belonging to identified hits (step 4). Reads are partitioned by genome assembly according to their strongest mapping. If a species with variants defined within the variant profiles file is detected during step 3, then the BAM file containing reads mapped to the specified reference fasta is passed to the variant profiling wing of Afanc `screen`. The reports from steps 3, 4, and variant profiling are collected and used to create a final report (step 5).

#### 2.3.1 Kraken 2 Report Disambiguation

Kraken 2 produces an extremely complex and potentially ambiguous metagenomic report, which details every possible match for the read set (see Figure 6). Interpreting this report can be extremely challenging, particularly in instances where the dataset is compound, consisting of multiple species and/or variants. This can be seen in figure 6, where every species which has at least 1 read attributed to it is reported. To solve this problem, a novel algorithm was developed to optimally identify the most likely species and variants present in a read set.

**Figure 6.**
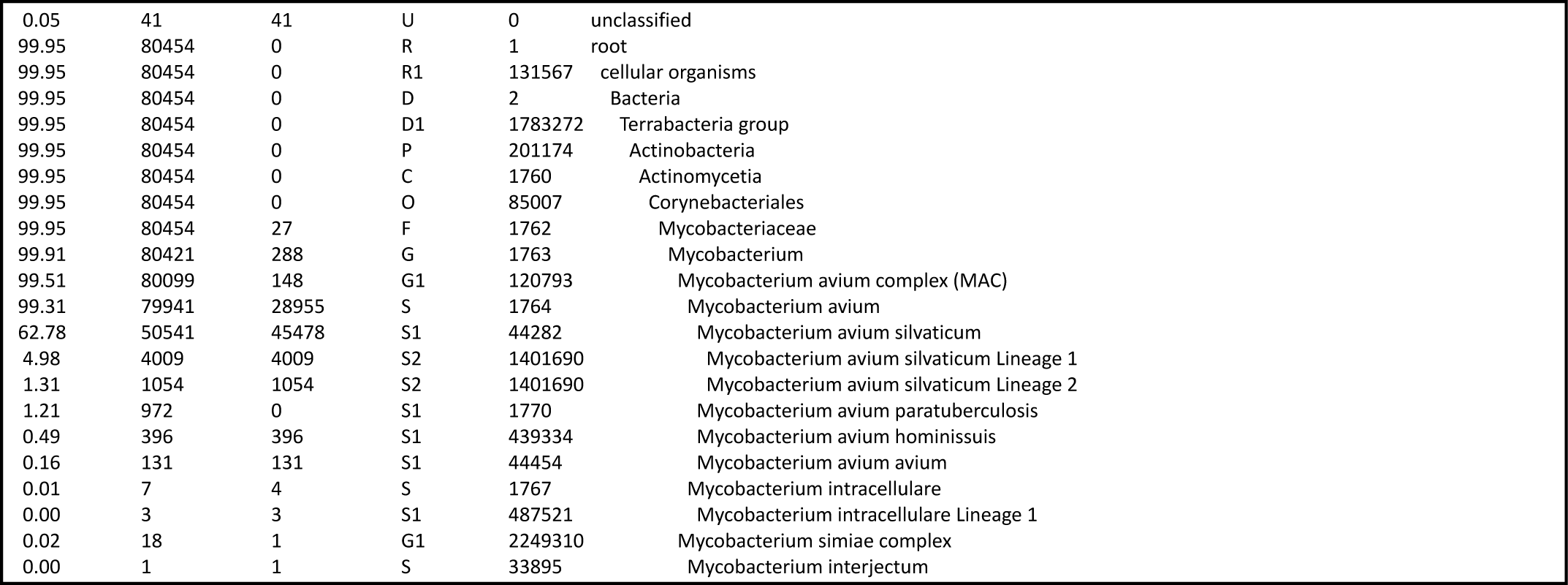
An example of a Kraken 2 report.

A Kraken 2 report is a hierarchical tree where nodes refer to individual taxa, topology is determined by taxonomic relations, and node weight is defined by the percentage of reads which were assigned to the taxon rooted at this node. Identifying key nodes within this tree must be carried out in a sensitive manner.

First-pass screening takes place by identifying species level or higher nodes which exceed a user defined global threshold (by default, this is 5.0%). This global threshold represents the minimum percentage of reads which must be attributed to a particular clade to consider it a putative hit. The branches rooted at each of these nodes are then traversed to find the maximally scoring tip nodes. A node is considered a hit if it exceeds a local elastic threshold (see Section 2.3.1.1). The tree is then subjected to the Bayesian Read Redistribution algorithm (see Section 2.3.1.2), and the scoring hit nodes identified in the previous step are reassessed to find the maximally scoring node.

##### 2.3.1.1 Elastic Threshold Calculation

The elastic threshold is calculated using the ANI values found within the variant index. The calculation is dependent on whether the node has a parent taxon which exists within the variant index, and therefore the ANI between the child and parent taxa exists.

Consider a node *n*∈*N* where *N* is the set of all species level or lower nodes which exceed the global threshold, and *p_n_* is the parent node of *n*. Given some similarity function *f* (which in this case, is the ANI), the ratio between the normalised parent and child ANI is

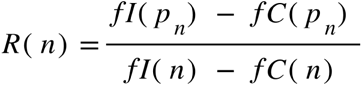

Where:

*fI*(*n*) = the mean intrataxon ANI of node *n*
*fC*(*n*) = the mean parent-child ANI of node *n*

The lower bound threshold weight for *n* is therefore

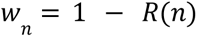

And the lower bound read count threshold for node *n* is therefore

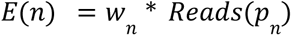

Where *Reads*(*p_n_*) is the number of reads assigned to the parent taxon of *n*.

##### 2.3.1.2 Bayesian Read Redistribution

When constructing a database using a large number of similar taxa, type I error (false-positive error) is very common during read assignment by Kraken2. This necessitates redistribution of reads between nodes within the Kraken2 report tree. This is achieved using a Bayesian approach, utilising the elastic threshold and the ANI between taxa rooted at that node.

Consider a tree *T* as a strict linearly ordered set of nodes, rooted at *T*_0_. A branch *b_n_* can be defined as a subtree of *T* rooted at node *n* ∈ *T*, such that *b* ⊆ *T*. Given a threshold *u* where 0 ≤ *u* ≤ 1, the set of all nodes which exceed their upper bound weighted elastic threshold is

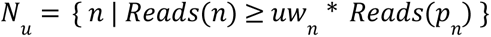

Symmetrically, given a threshold *l* where 0 ≤ *l* < *u*, the set of all nodes which fall below their lower bound weighted elastic threshold is

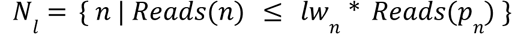

Reads are commuted from nodes in *N_l_* to nodes in *N* if they satisfy two criteria:

1) *N_i_* and *N_u_* are rooted at the same parent node.
2) *N_l_*and *N_u_*are the same taxonomic level.

For example, consider the tree in Figure 7. Reads can be commuted between all taxa on level 1 (e.g from M.bovis to M.tb L1/L2) since they share the same parent, but not between taxa on level 2 (e.g. M.b BCG to M. tb L2.1) since they do not share the same parent, and are cousin taxa. However, reads which are redistributed from M.bovis to M.tb L2 are trickled into L2.1 in a number which conserves the proportion of reads from M.tb L2 which were assigned to subtaxa L2.1 prior to commuting. Consequently, monotypic tip level taxa which do not exceed the elastic threshold prior to commuting cannot exceed it after commuting, even if the parent does.

**Figure 7.**
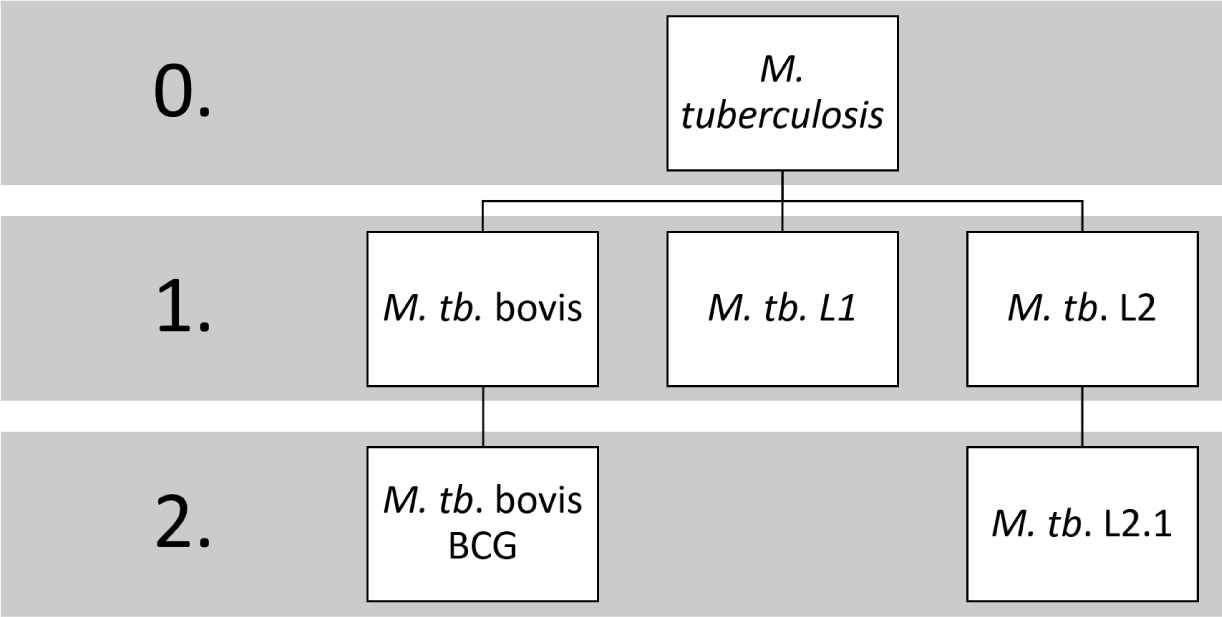
A hierarchical tree representing a subset of M. tuberculosis taxonomy.

Reads are commuted between taxa in an entirely probabilistic manner, whereby the number of reads commuted to taxon *T_i_* from taxon *T_j_* is determined by their ANI and the percentage of reads assigned to the shared parent taxon which were further assigned to *T*_i_.

Given a tree *T* where *T*_*i*_ refers to node *i* within the tree, and a set of reads *R*. The elastic threshold of *T_i_* is *E*(*T_i_*), and *Reads*(*T_i_*) ⊆ *R* is the set of all reads assigned to *T_i_*. The set of all misassigned reads *R_m_* is considered as the set of all reads assigned to nodes which fall below the lower bound weighted elastic threshold

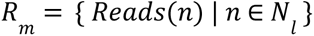

The probability of a read *r* ∈ *R* being misassigned is therefore

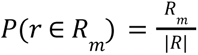

The ANI between the sequences of nodes *T_i_* and *T_j_* is *f*(*T_i_ T_j_*). The probability of a read belonging to sequence *T_i_* is

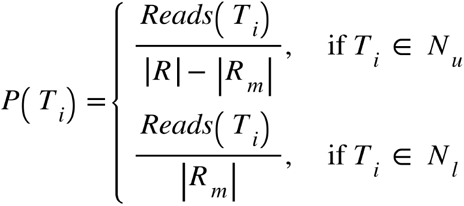

Finally, the probability that a read *r* ∈ *R_m_* misassigned to taxon at node *T_i_* belongs to the taxon at node *T_j_* is

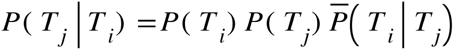

Where

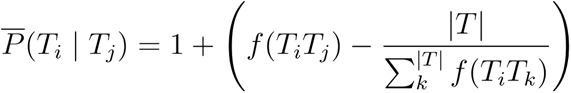

##### 2.3.1.4 Variant Profiling

Variant profiling is achieved by taking a set of reads mapped to a reference genome assembly (in BAM format) and a set of variant definitions (in BED format) and querying the mapping BAM file for mutations found within the variant definitions file. Variants are determined to be present if all variant alleles (mutations) defined within the bed file for a variant are detected with a minimum coverage of 5x and pass a probability threshold. Variant allele probability is calculated as *P*(*v*) = *d*/*v* where *d* is the depth of coverage at this position, and *v* is the variant allele count at this position. The probability threshold is calculated as 0.15 standard deviations from the mean variant allele probability across the set of all alleles used to define each variant.

## 3. Method

### 3.1 Databases

The `autodatabase` module was used to construct a database from an extensive repertoire of Mycobacteriaceae genome assemblies available on GenBank (n=223 across 139 species). The majority of species (n=133) were represented by single genome assemblies. Select species, for which variant level identification was of greater importance (n=6), were represented by multiple genome assemblies of subspecies and variants. This database includes a collection of discrete *Mycobacterium tuberculosis* lineages (covering lineages 1, 2, 3, 4, 5, 6, 7, 9, the majority of their sublineages down to 4th order, and bovis, caprae, and orgis). A full list of genome assemblies used to construct the Mycobacteriaceae database can be found in Supplementary Materials.

A variant catalogue was used to define lineages within *Mycobacterium tuberculosis*. Variant profiles from Napier *et al*. (2020) were used to construct the variant catalogue, with some adjustments made to fine tune the depth of sub-lineage definitions within each lineage. This variant catalogue was used in the variant profiling wing of Afanc `screen`.

### 3.2 Simulated Data

Reads were simulated using genome assemblies from the family Mycobacteriaceae using ART (Huang et al., 2011). Three Mycobacteriaceae datasets were used. Dataset M1 consists of 846 read sets simulated from 141 separate Non-Tuberculosis mycobacteria (NTM) species (n=136) and subspecies (n=5) (see Table S1 in the supplementary materials). Each species is represented by 6 read sets covering 1%, 5%, 10%, 20%, 50% and 100% of the genome covered to 10x. Dataset M2 consists of a further 6 read sets, constructed from 8 key species and variants (*Mycobacterium kansasii*, *Mycolicibacterium fortuitum*, *Mycobacterium interjectum*, *Mycobacteroides chelonae*, *Mycobacterium intracellulare* subsp. *chimaera*, *Mycobacterium avium* subsp. *paratuberculosis*, and *Mycobacterium tuberculosis* variant *bovis* BCG), resulting in complex compound datasets. These were simulated using 5%, 20%, 40%, 60%, 80% and 100% of each constituent genome, covered to a depth of 40x (see Table S2 in the supplementary materials).

### 3.3 *Mycobacterium tuberculosis* Lineage Profiling

401 *Mycobacterium tuberculosis* paired-end FASTQ readsets, covering all characterised *M. tuberculosis* lineages, many of their sublineages, and variants (bovis, caprae, and orgis) were used to test the variant profiling wing of Afanc screen. A list of accessions and their reported lineage designation can be found in Table S3 in the supplementary material.

### 3.4 Benchmarking

Afanc was benchmarked against Mykrobe v0.12.1, Kraken v2.1.2, Bracken v2.7 and KrakenUniq v0.5.8 using datasets M1-3. For *M. tuberculosis* lineage profiling, Afanc was benchmarked against Mykrobe v0.12.1 and TB-profiler v4.2.0. The standard Mykrobe and TB-profiler databases were used. The variant profile used with Afanc was modified from the standard TB-profiler database to remove instances where sub-lineages were reported in the absence of SNPs present in their parental lineages.

## 4. Results

The results from running Afanc, Mykrobe, Kraken, Bracken, and KrakenUniq on dataset M1 can be seen in Figure 8. Afanc reported the correct species or subspecies in all datasets across all coverage cohorts. Mykrobe fails to identify any species/subspecies correctly when 1% of the genome is covered, and reports only 2 species correctly (*Mycobacterium heidelbergense* & *Mycobacterium peregrinum*) when 5% of the genome is covered. However, when at least 10% of the genome is covered, Mykrobe performs substantially better, reporting correctly on 71-86% of NTMs. Bracken consistently outperformed KrakenUniq across all cohorts, and there was little difference in the pass rate across each cohort. Bracken reported the correct species in 85% of cases across all coverage cohorts. Likewise, KrakenUniq reported the correct species in 53% of cases across all coverage cohorts. Bracken has the highest rate of underqualifications across the 5 subspecies. Neither Afanc nor Mykrobe underqualified any subspecies.

**Figure 8.**
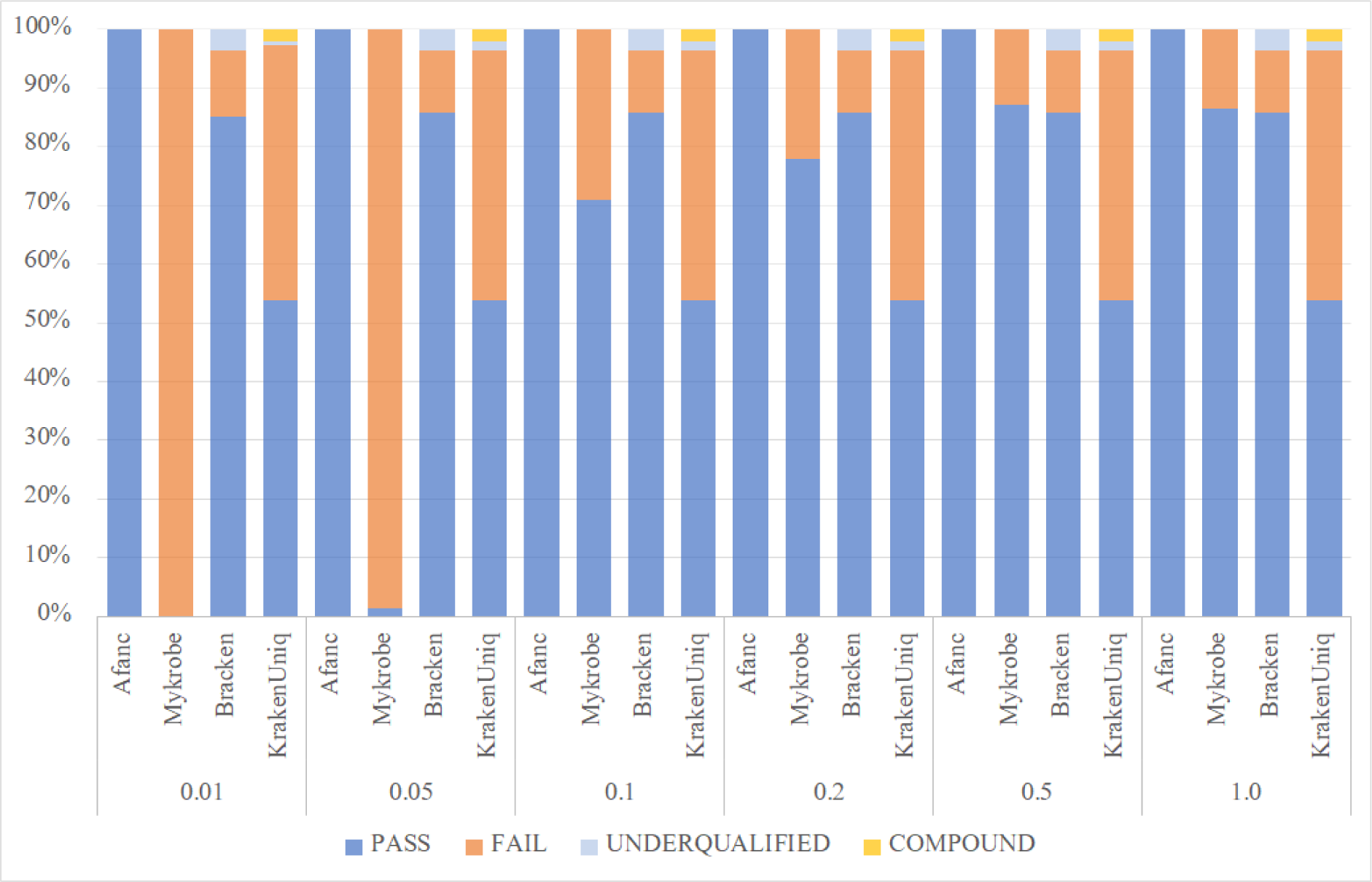
The results of running Afanc, Mykrobe, and TBprofiler using dataset M1. PASS = exact species/subspecies match; FAIL = reported species is incorrect. UNDERQUALIFIED = reported species is a parent of Truth; COMPOUND = multiple species/variants reported including Truth.

The results of running all tools on dataset M2, containing complex compound datasets with a diverse number of Mycobacteriaceae species and variants can be found in Figure 9. Afanc identified all members of the compound dataset correctly across all datasets. The Afanc variant profiling module successfully identified *M. tuberculosis bovis* BCG to sub lineage level (BCG La1.2) in this dataset down to 20% coverage across the sample genome to a depth of 40x. At 5% genome coverage, the coverage across the *M. tuberculosis* var. *bovis* BCG genome was too low to capture all SNPs necessary to positively identify it. However, Afanc identified BCG within this dataset during the initial screening step. Mykrobe failed to elucidate each species robustly within this dataset, particularly where only a fragment of the genome was present. At 100% coverage across each genome (M2.6), Mykrobe was able to identify *M. chelonae*, *M. kansasii*, *M. fortuitum*, *M. chimaera*, *M. avium*, and *M. tuberculosis bovis* BCG correctly. Bracken consistently identified *M. intracellulare*, *M. intracellulare chimaera*, and *M. chelonae* across all datasets. However, it fails to elucidate all constituent species in each dataset. Furthermore, Bracken erroneously reported the presence of *M. haemophilum* and *M. leprae*. KrakenUniq identifies *Mycobacterium intracellulare*, *Mycobacterium intracellulare chimaera*, *Mycobacterium avium*, *Mycobacterium kansasii*, *Mycolicibacterium fortuitum*, and *Mycobacteroides chelonae* in all datasets, but fails to report *M. interjectum* or *M. tuberculosis bovis* BCG.

**Figure 9.**
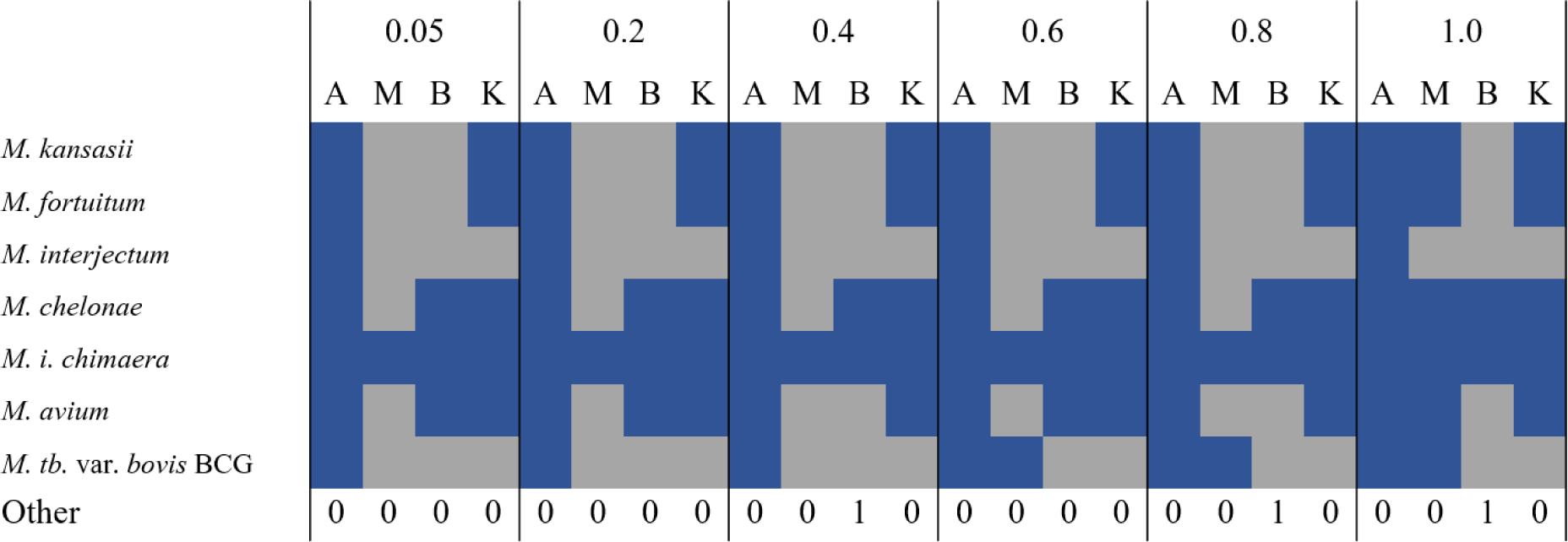
The results of running Afanc, Mykrobe, and TBprofiler using dataset M2 across all coverage cohorts. A = Afanc, M = Mykrobe, B = Bracken, K = KrakenUniq. Blue = species/subspecies/variant reported. Grey = species/subspecies/variant not reported. Other = the number of un-listed Mycobacteriaceae reported.

The results of running Afanc, Mykrobe and TBprofiler on the *M. tuberculosis* lineage dataset can be seen in Figure 10 and Table S4. These results indicate a high level of similarity between the lineages reported by Afanc and TBprofiler, with pass rates of 0.925 and 0.895 respectively. Mykrobe has a pass rate of 0.693. Each profiler has a very similar rate of overqualification (0.023-0.038). Overqualifications seem to be very consistent between Afanc and TBprofiler, whereby overqualified lineages are often reported identically. The results from lineage 4 indicate a difference in categorisation between Afanc, TBprofiler, and Mykrobe. Lineages 4.7, 4.8, and 4.9 are reported by Afanc and TBprofiler, however, Mykrobe subsumes these into a compound lineage 4.10. Consequently, Mykrobe cannot distinguish these lineages from each other. Similarly, for lineages 3, 5, and 6, Mykrobe does not perform lower level lineage disambiguation in many cases. Some sub-lineages of 3.1 are reported, but 3.1 and 3.1.3 are classified as lineage 3. Lineages 5 and 6 are not further classified into sub-lineages. The analysis also indicates that the basis for the definition of Lineage 5 is problematic, as all profilers either underqualify or overqualify a substantial portion of it. TB-profiler appears to incorrectly classify lineage 3.2 as lineage 3.1.3, despite the SNPs for parental lineage 3.1 not being present in these samples.

**Figure 10.**
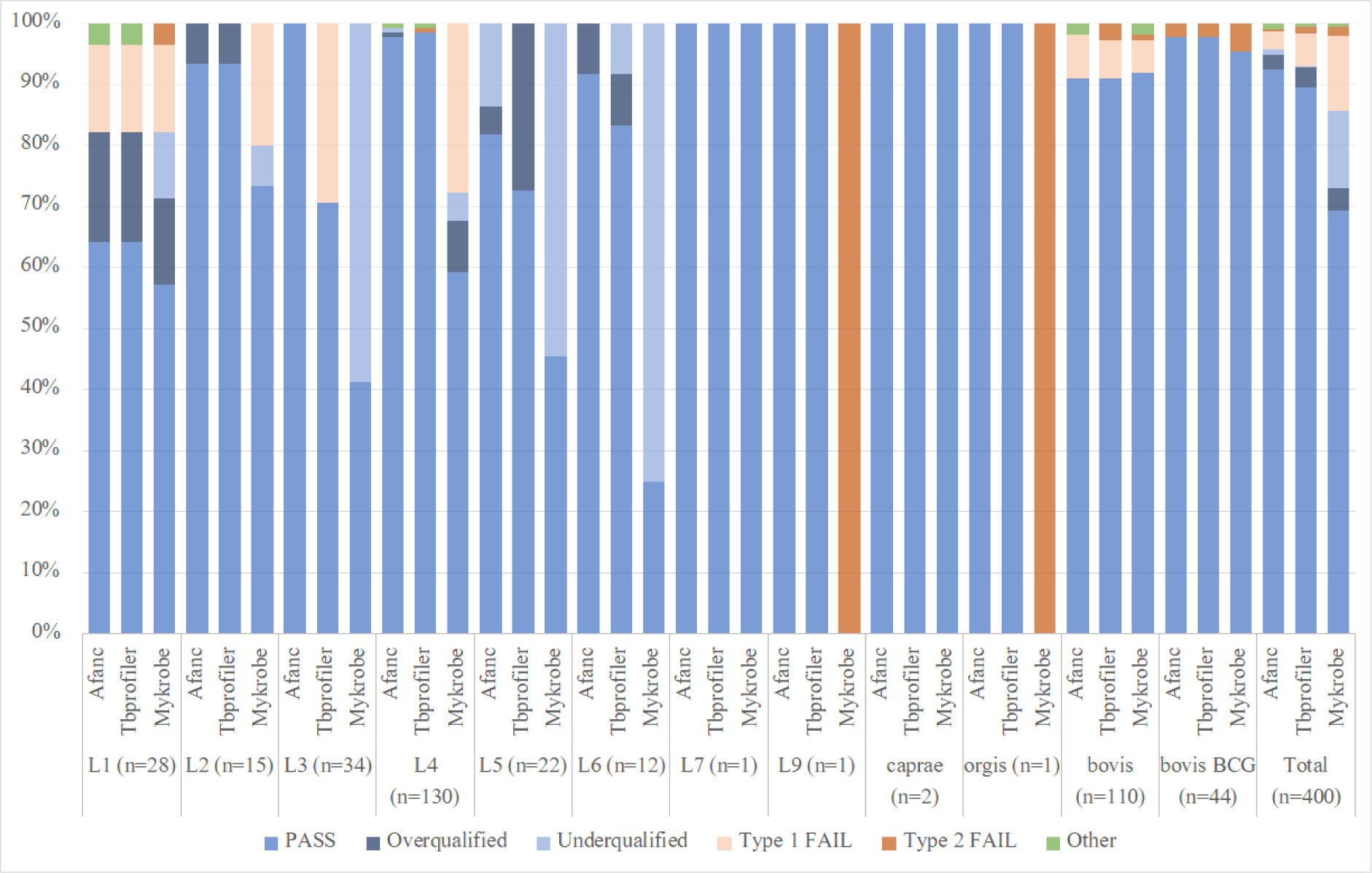
The results of running Afanc, Mykrobe, and TBprofiler using the lineage dataset. PASS = exact lineage match; Overqualified = reported lineage is a sub-lineage of Truth; Underqualified = reported lineage is a parent lineage of Truth; Type 1 FAIL = reported lineage is a cousin lineage to the Truth; Type 2 FAIL = reported lineage is incorrect; Other = run fail or indication of low coverage.

## 5. Discussion

The challenges associated with extracting DNA from clinical Mycobacteriaceae samples often means that coverage is low and fragmented. Mycobacteriaceae may also be relatively slow growing, meaning that it is imperative that speciation tools for clinical use are able to operate with samples that are of variable quality. The provision of tools that enable reliable identification of Mycobacteriaceae and other pathogen species where there is poor biological signal is therefore essential for the development of clinical and public health genomic services. The comparison of speciation tools tested against dataset M1 demonstrate that the screening algorithm used by Afanc is more powerful both in the identification of NTM species and subspecies than similar approaches used by Bracken and KrakenUniq, and the probe-driven approach used by Mykrobe. Afanc is able to correctly and reliably identify NTMs where only small fragments of the genome are present, down to 1% breadth of coverage, far beyond the limitations of currently available cutting edge tools.

Clinical datasets can consist of multiple species and sub-species populations, enormously increasing the complexity of sample profiling. Results from processing dataset M2 highlights the difficulties associated with *in silico* metagenomic profiling of compound datasets. Afanc performs extremely favourably when compared to other metagenomic profilers across all coverage cohorts, with only KrakenUniq also performing consistently well across all cohorts. However, KrakenUniq required an *ad hoc* 5% threshold applied to the report to filter out large numbers of false-positive taxa. This also resulted in significant over correction, whereby taxa at a level lower than species, which will have very few reads assigned uniquely to them, were incorrectly filtered out. In the case of KrakenUniq, this resulted in the loss of M. tuberculosis var. bovis BCG within its report, which is a major issue that would preclude the use of this software in a clinical or public health setting. The same is true for Bracken, but with worse overall species detection. Mykrobe consistently and correctly reported the presence of *M. intracellulare chimaera* (reported as *M. chimaera* in accordance with previous taxonomic nomenclature) across all coverage cohorts, and *M. tuberculosis bovis* var. BCG in cohorts where total proportional genomic coverage was 0.8 and 1.0, but failed to report all other species across all cohorts. This dataset highlights the advantages of Afanc in dealing with compound datasets compared to the other tools listed. In particular, the strength of the dual screening methodology (first pass metagenomic screening followed by variant profiling) employed is demonstrated in instances where the biological signal for a species or variant is too low to positively identify all variant defining polymorphisms.

Afanc compares favourably with both TB-profiler and Mykrobe when carrying out TB lineage profiling, with the highest proportion of correctly profiled lineages (0.925), and the lowest type-2 failure rate (0.003). It has a higher underqualification rate (0.01) than TB-profiler (0.003). This is a result of the removal of ambiguity in lineage 5 within the SNP profile. Lineages which were previously classified within the *Mycobacterium africanum* species (5 & 6) are poorly characterised in comparison to other TB lineages, and consequently a conservative approach to reporting these lineages was taken. The results indicate that Mykrobe adopts a similar approach for some lineages. Results of TB lineage profiling from Afanc and TB-profiler have a high degree of concordance. This is undoubtedly due to the fact that they both use a SNP profiling approach, and used a similar set of lineage SNP profiles. All tools perform very similarly on bovis and bovis BCG datasets, with Mykrobe exhibiting a slightly higher Type 2 failure rate when processing BCG samples. There are a number of instances where all three profilers report BCG from datasets which ostensibly belong to bovis, and one instance where Lineage 1.2.2 was concurrently reported by all profilers when processing a BCG dataset. It is likely that this is as a result of datasets being mislabelled on GenBank.

Afanc has been demonstrated to outperform the cutting edge speciations tools at species/subspecies level characterisation, in disambiguating compound samples, and for lineage level disambiguation of Mycobacteriaceae across all testing parameters. Afanc also allows the user to construct their own database from a bespoke set of species and variants, or utilise their own set of variant definitions, ensuring that Afanc can be used for the analysis of any pathogen species, and providing a system for the construction of databases that keep up with the generation of new data. This also ensures that as novel species are defined, and new variants characterised, Afanc can be used to identify them.

Afanc is designed to be run on unix systems via the command line. This allows for seamless integration into bioinformatics workflows and pipelines. Due to the low computational requirements of Afanc, it can be run on a personal computer, obviating the need for HPC or cloud computing platforms. Installation instructions and a list of dependencies are detailed on the github page, which can be found in the Software Availability section of this paper.

Currently, variant profiles must consist of single nucleotide polymorphisms (SNPs). In the future, we intend to expand the functionality of the variant profiler to support other classes of variants.

## 6. Conclusion

The accurate and reliable species and variant level identification of pathogens within clinical samples is a cornerstone of the work undertaken by medical and public health laboratories. Increasingly, it is also being recognised that tools that provide speciation, must also provide mechanisms to enable the updating and creation of higher quality databases than can be managed by a simple bulk download from the NCBI or EBI. Development of reliable *in silico* bioinformatics approaches and tools to enable the construction of better quality databases, combined with tools to exploit them is therefore of utmost importance. In a thorough analysis of the taxonomic landscape of the Mycobacteriaceae, and the lineages of *M. tuberculosis*, we have demonstrated Afanc to be a robust and highly sensitive tool in performing species, subspecies, and lineage level profiling in even the most complex and low signal multi-species datasets. Afanc outperforms the major contemporary cutting edge Mycobacteriaceae profilers currently available across all tested fields, allowing for more precise and reliable disambiguation. Furthermore, Afanc is an entirely general and species agnostic profiler, allowing the user to construct bespoke databases and provide their own set of variant definitions, thereby aiding in the future surveillance of pathogens of clinical importance, both extant and emerging. It is our hope that Afanc will be employed by medical and public health laboratories to form the backbone of speciation and variant characterisation workflows when dealing with clinical pathogen NGS data, and that researchers will find its autodatabasing capability to be of significant utility, and enabling better, more accurate speciation for a wide range of species and situations.

## Software Availability

Afanc is available at https://github.com/ArthurVM/Afanc.

## Author Contributions

T.C., A.M. and A.P. conceived of presented software. A.P. designed and wrote the first version of autoDatabase in nextflow. A.M. designed and wrote Afanc, carried out testing, and wrote the first draft of the manuscript. All authors contributed to editing and adjusting the final manuscript.

## Correspondence

Correspondence should be directed to A.M.

## Acknowledgements

I am grateful to Owen Jones and Tom Whalley, who provided advice and support when carrying out the work presented in this paper.

## Appendix Testing

All testing was carried out on a laptop running Ubuntu v20.04.4 LTS (Focal Fossa), with 32Gb RAM, and an intel i7 6 core CPU.

### Supplementary Materials

**Table S1.**
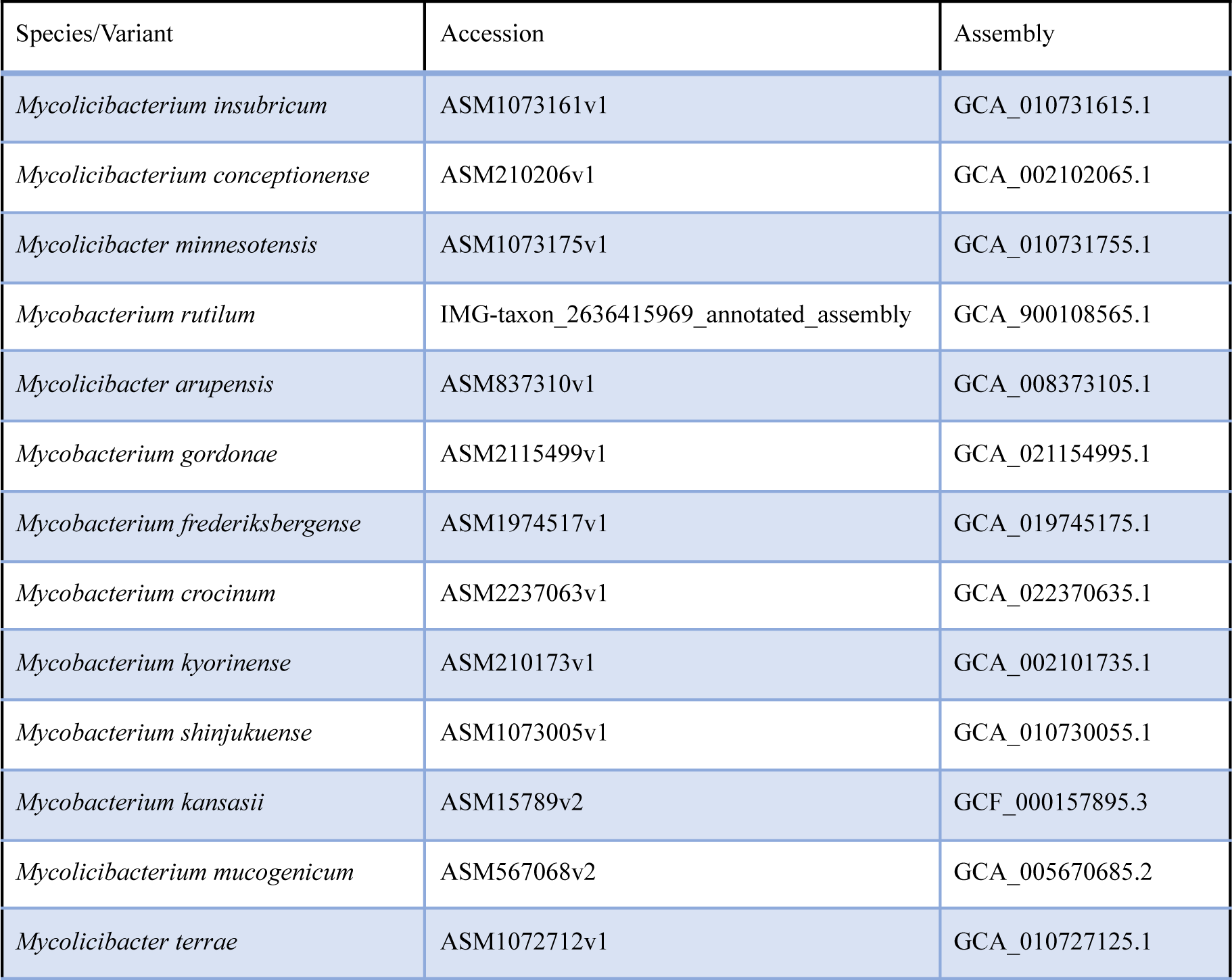
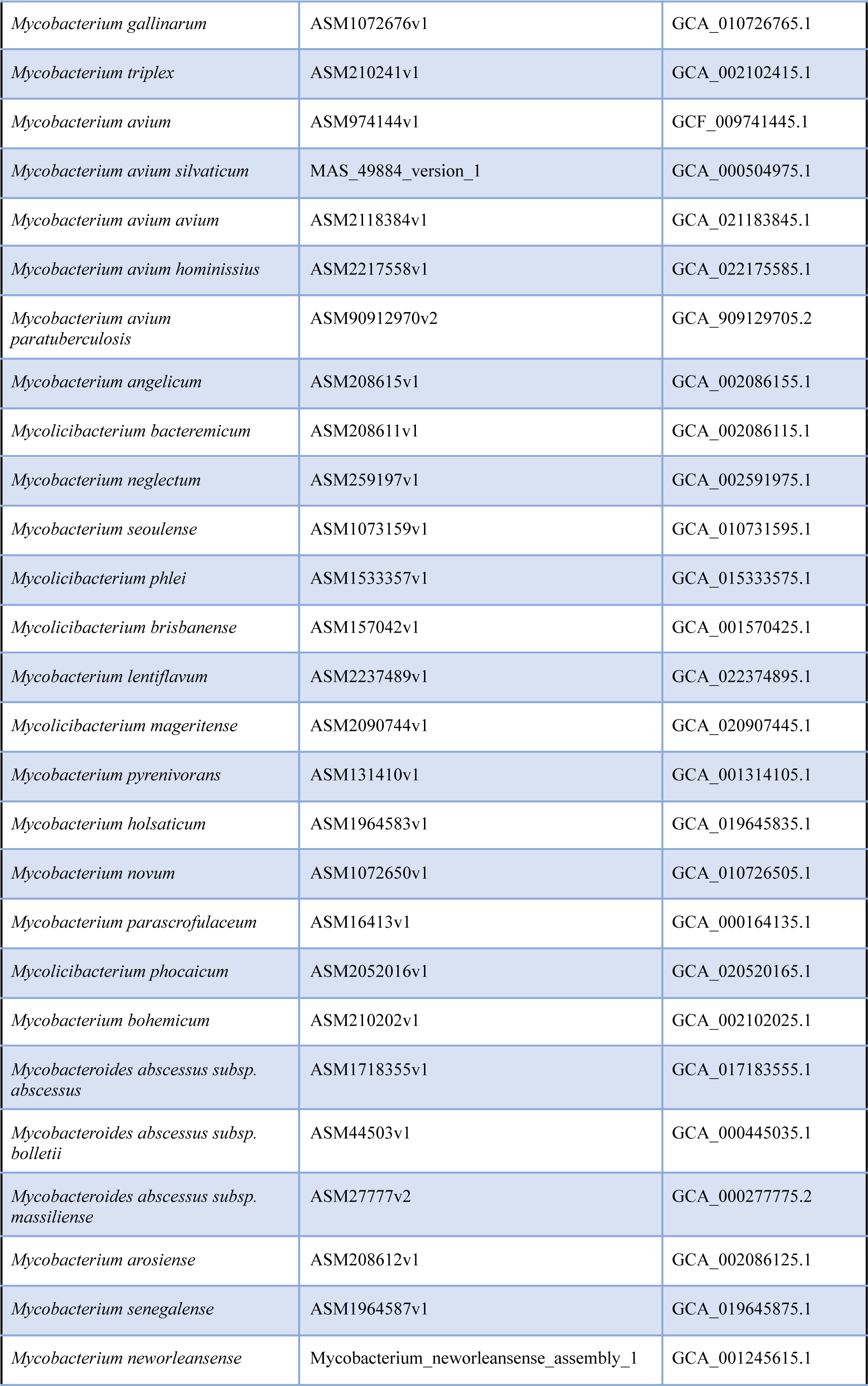
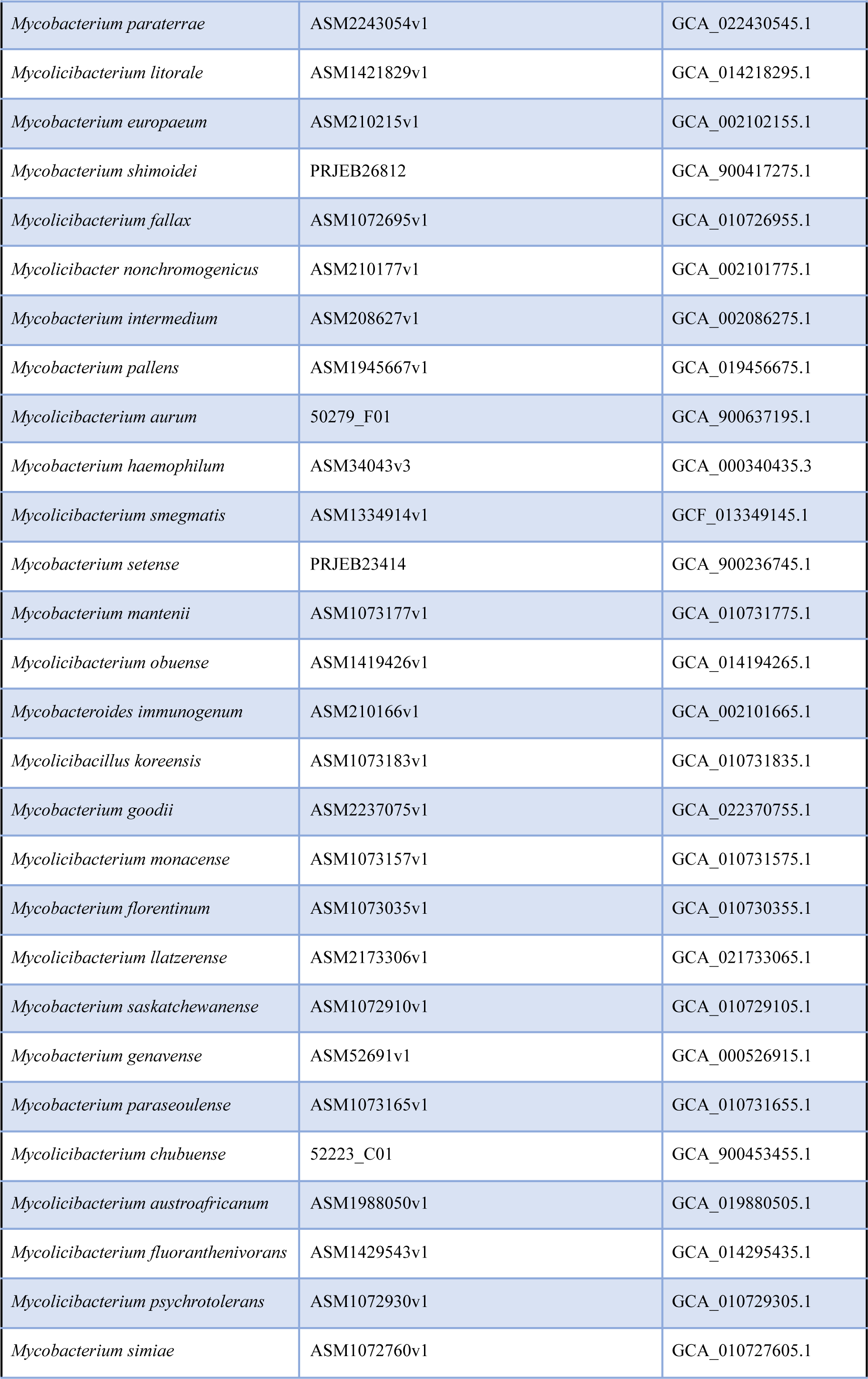
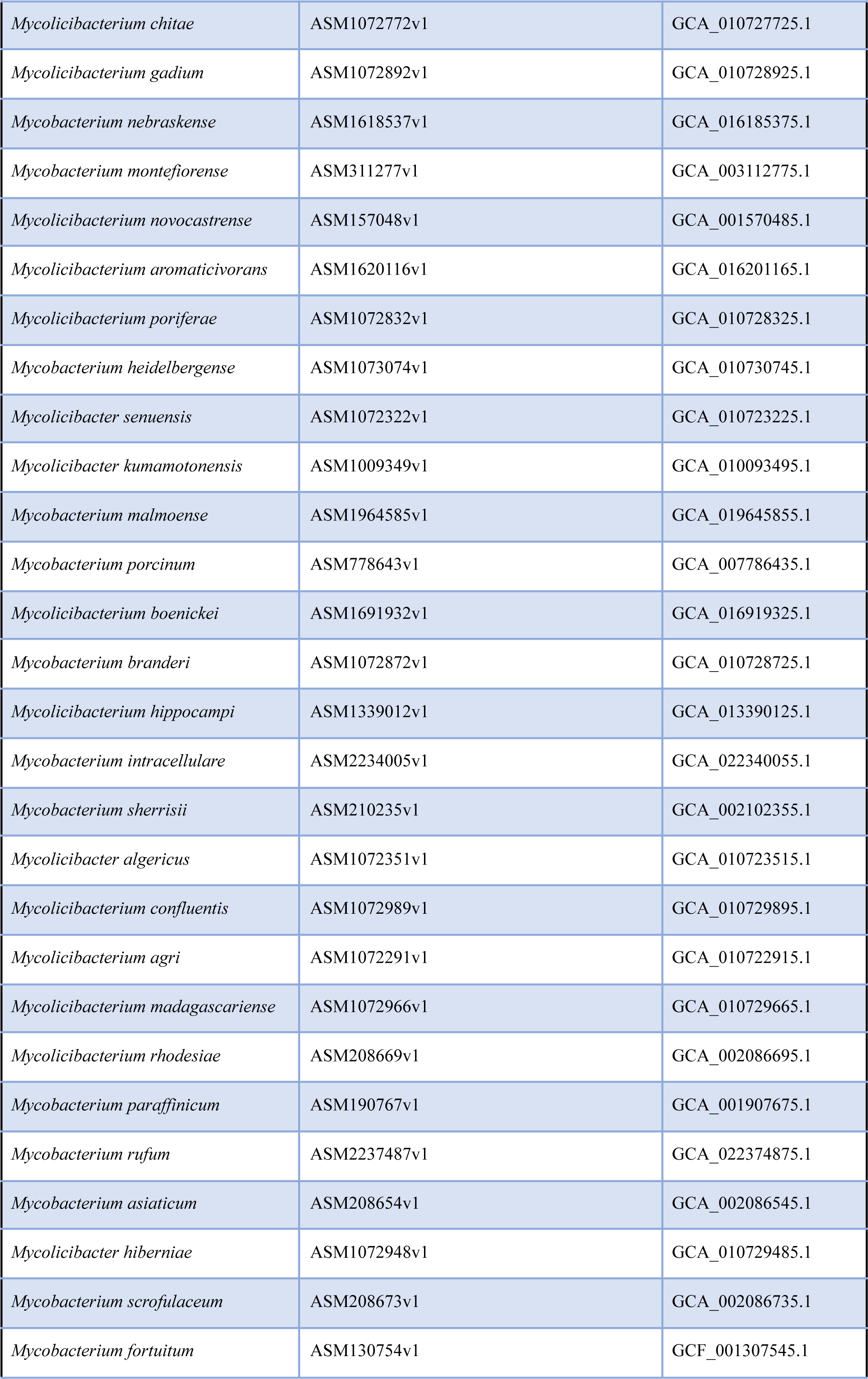
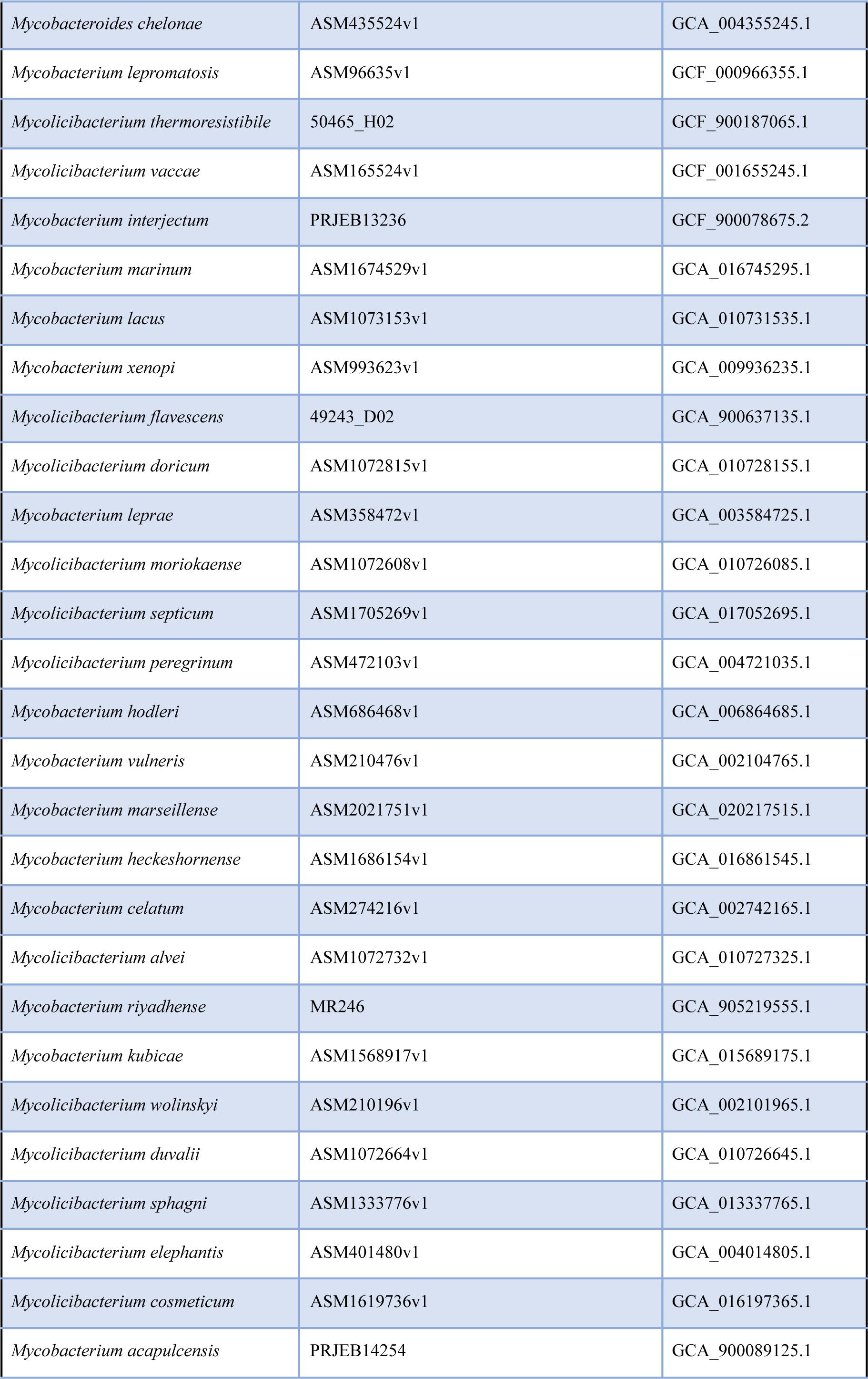
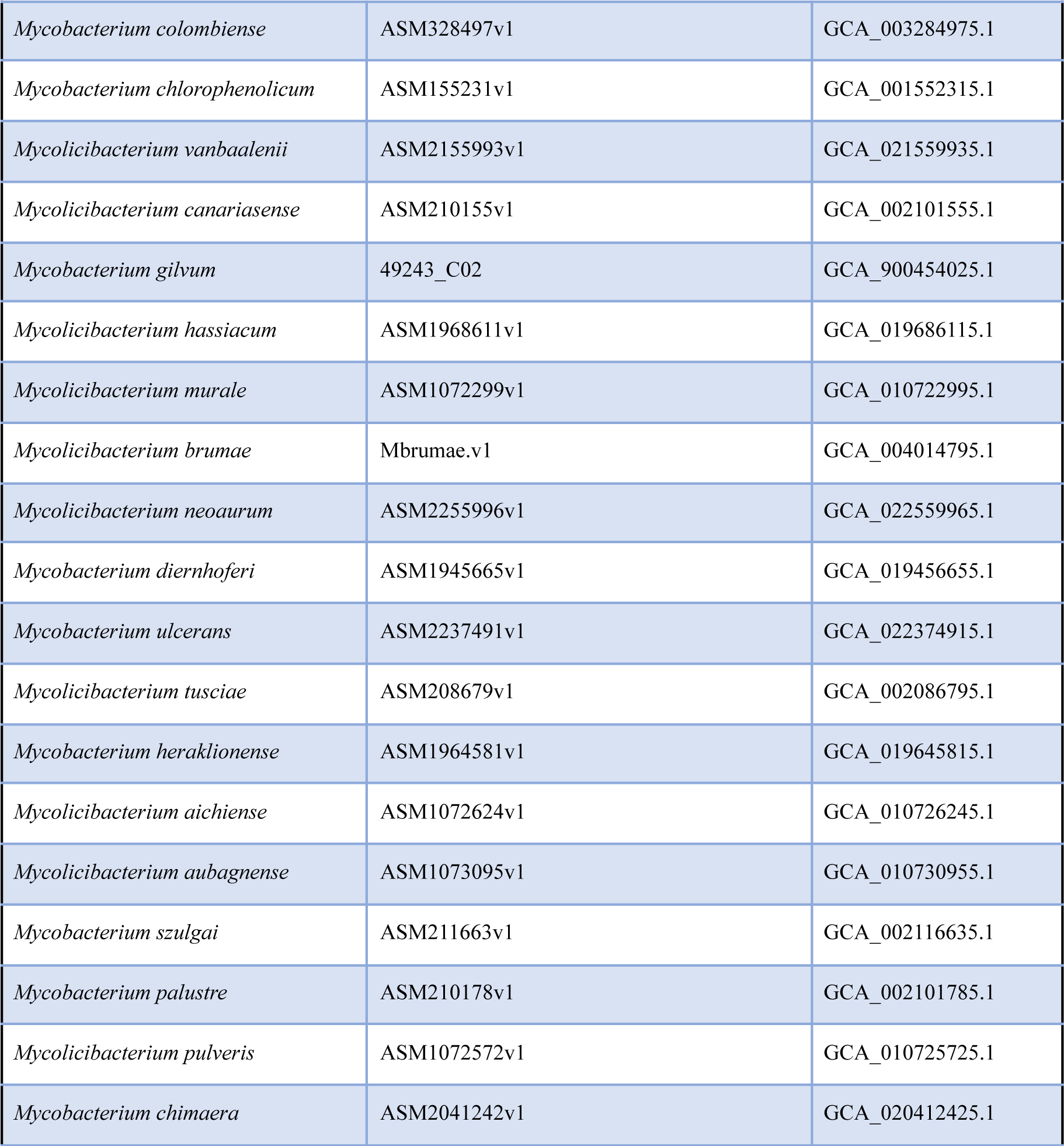
Dataset M1.

**Table S2.**
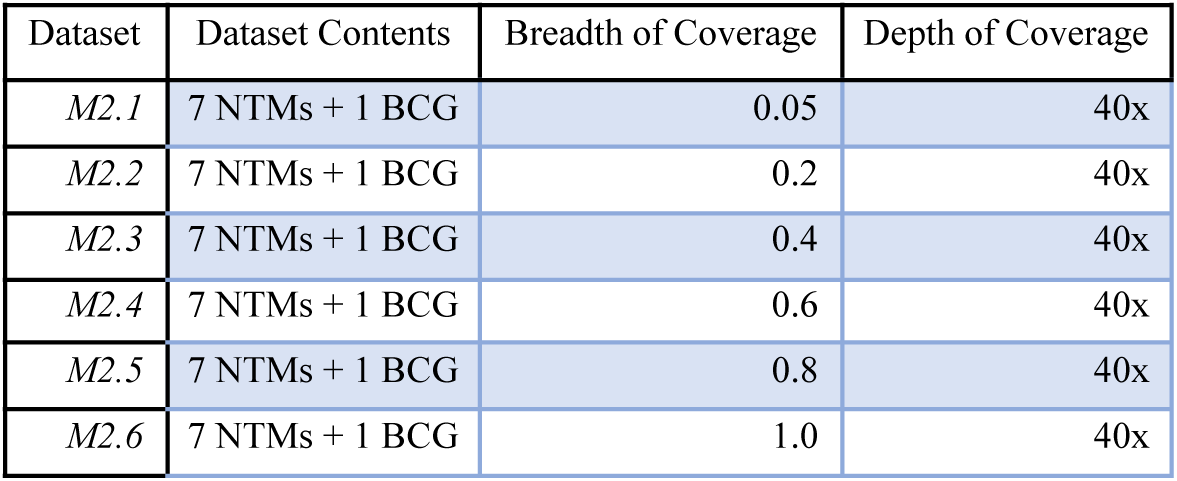
Dataset M2.

| Dataset | Dataset Contents | Breadth of Coverage | Depth of Coverage |
| --- | --- | --- | --- |
| M2.1 | 7 NTMs + 1 BCG | 0.05 | 40x |
| M2.2 | 7 NTMs + 1 BCG | 0.2 | 40x |
| M2.3 | 7 NTMs + 1 BCG | 0.4 | 40x |
| M2.4 | 7 NTMs + 1 BCG | 0.6 | 40x |
| M2.5 | 7 NTMs + 1 BCG | 0.8 | 40x |
| M2.6 | 7 NTMs + 1 BCG | 1.0 | 40x |

**Table S3.**
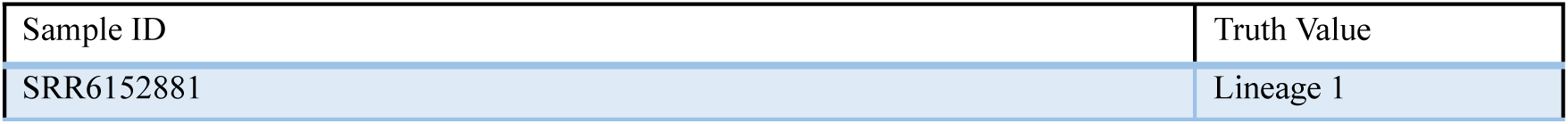
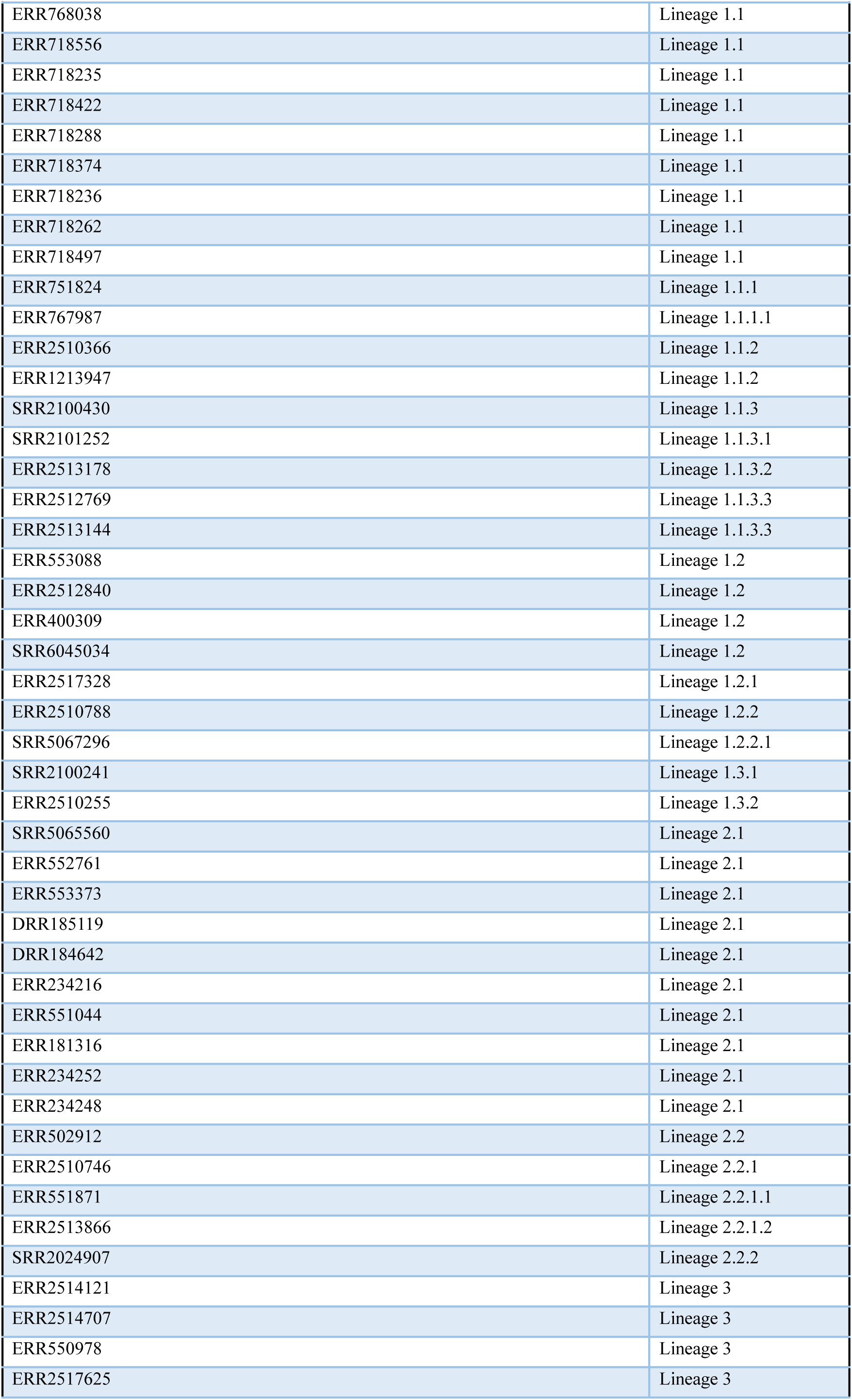
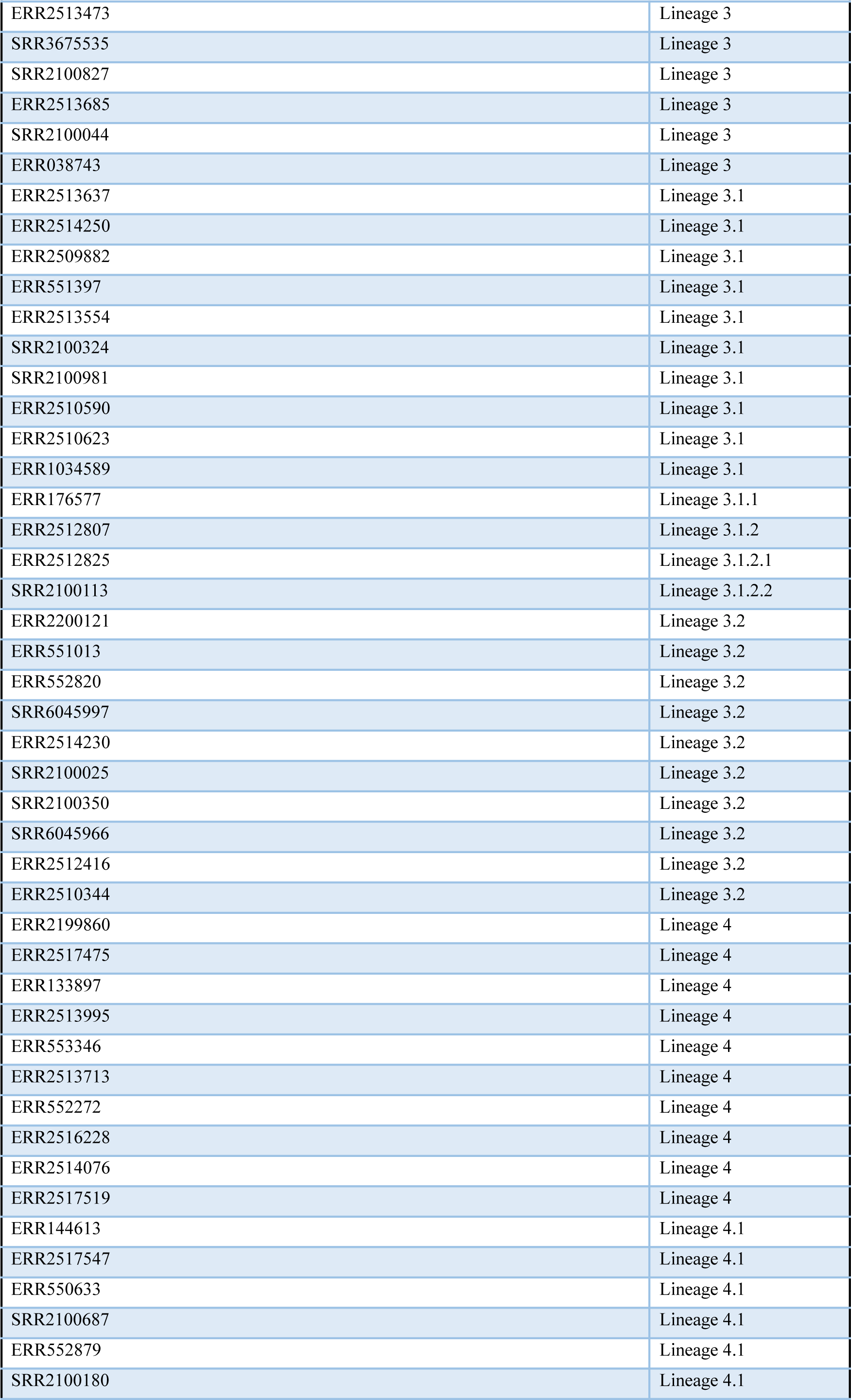
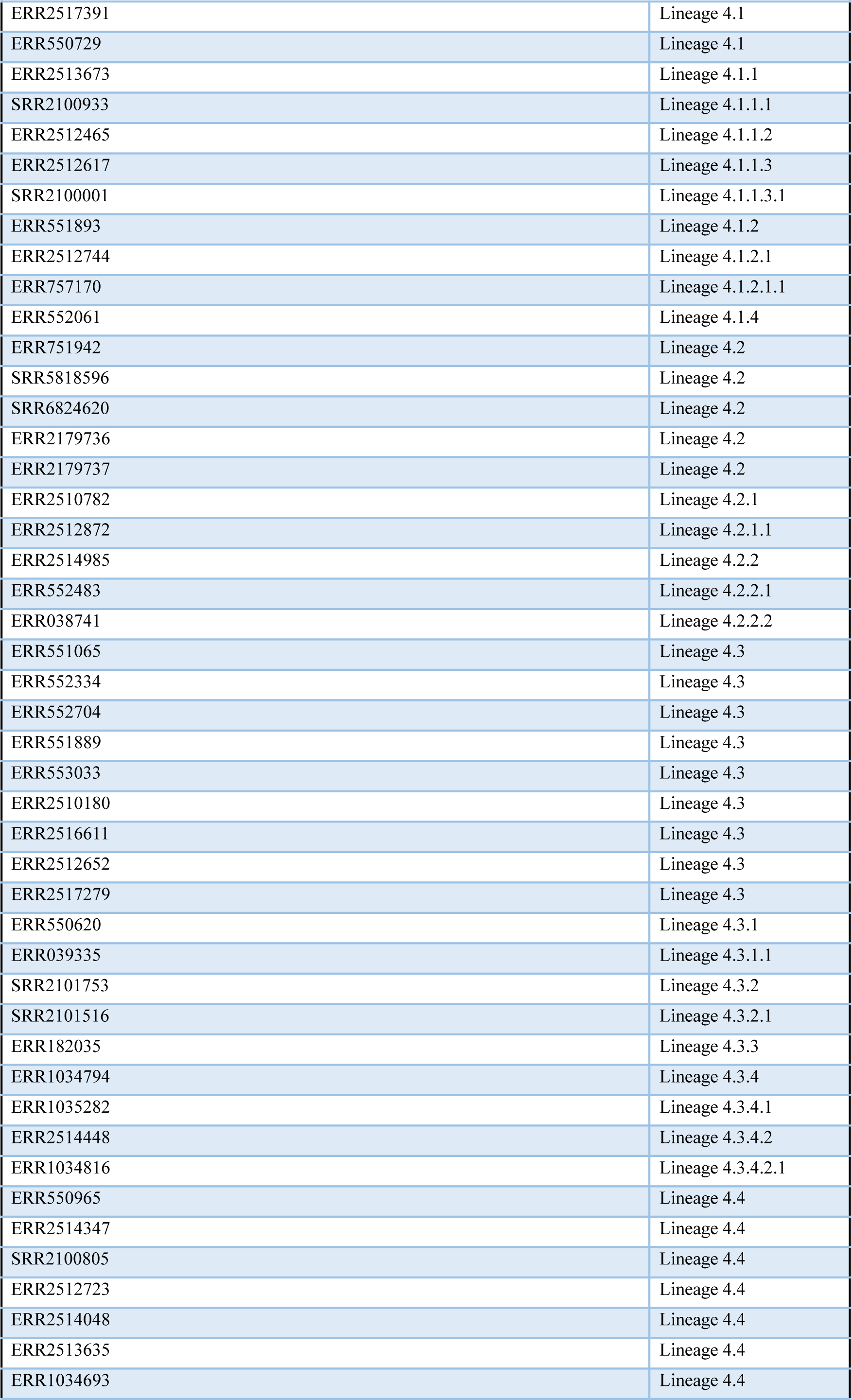
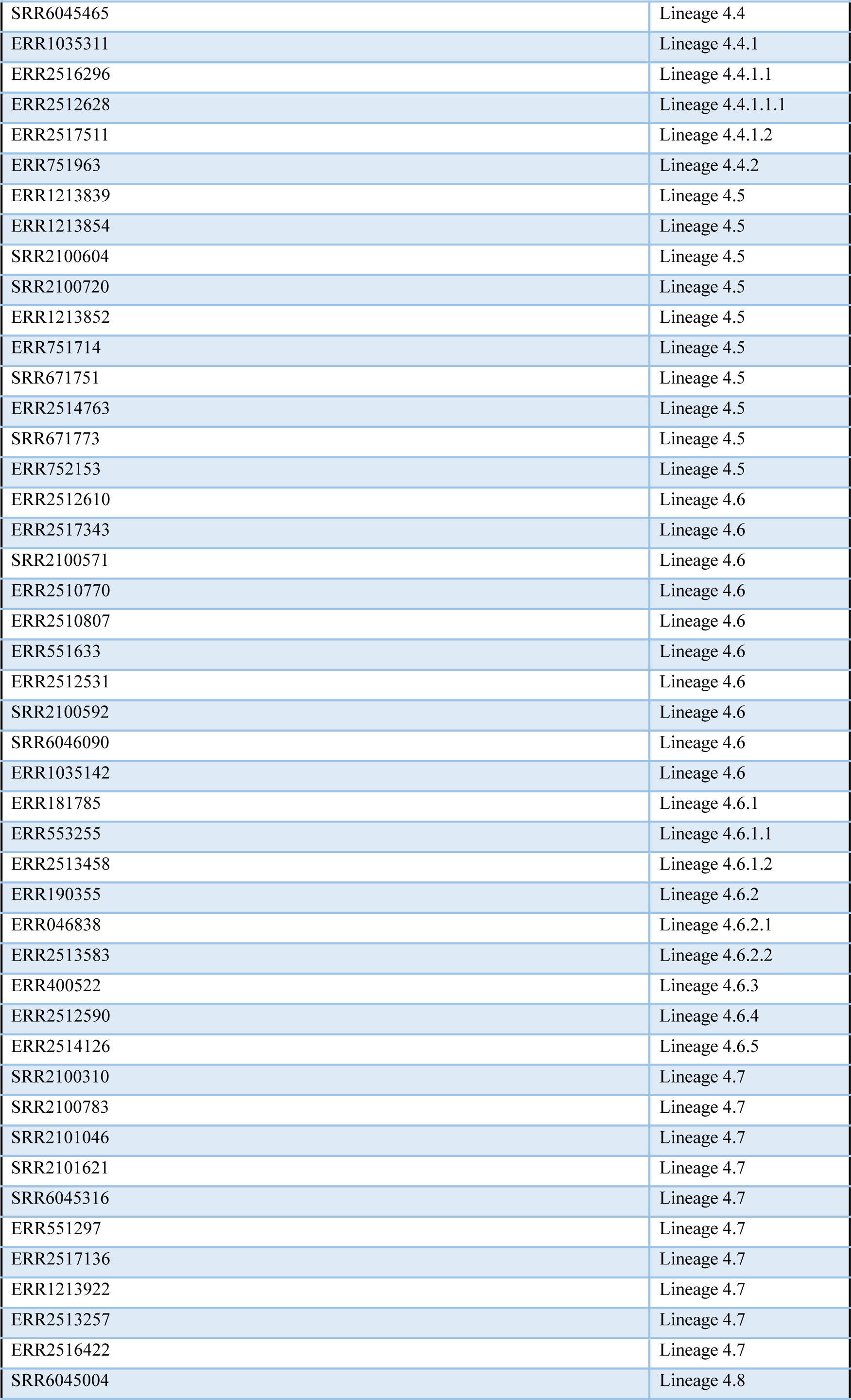
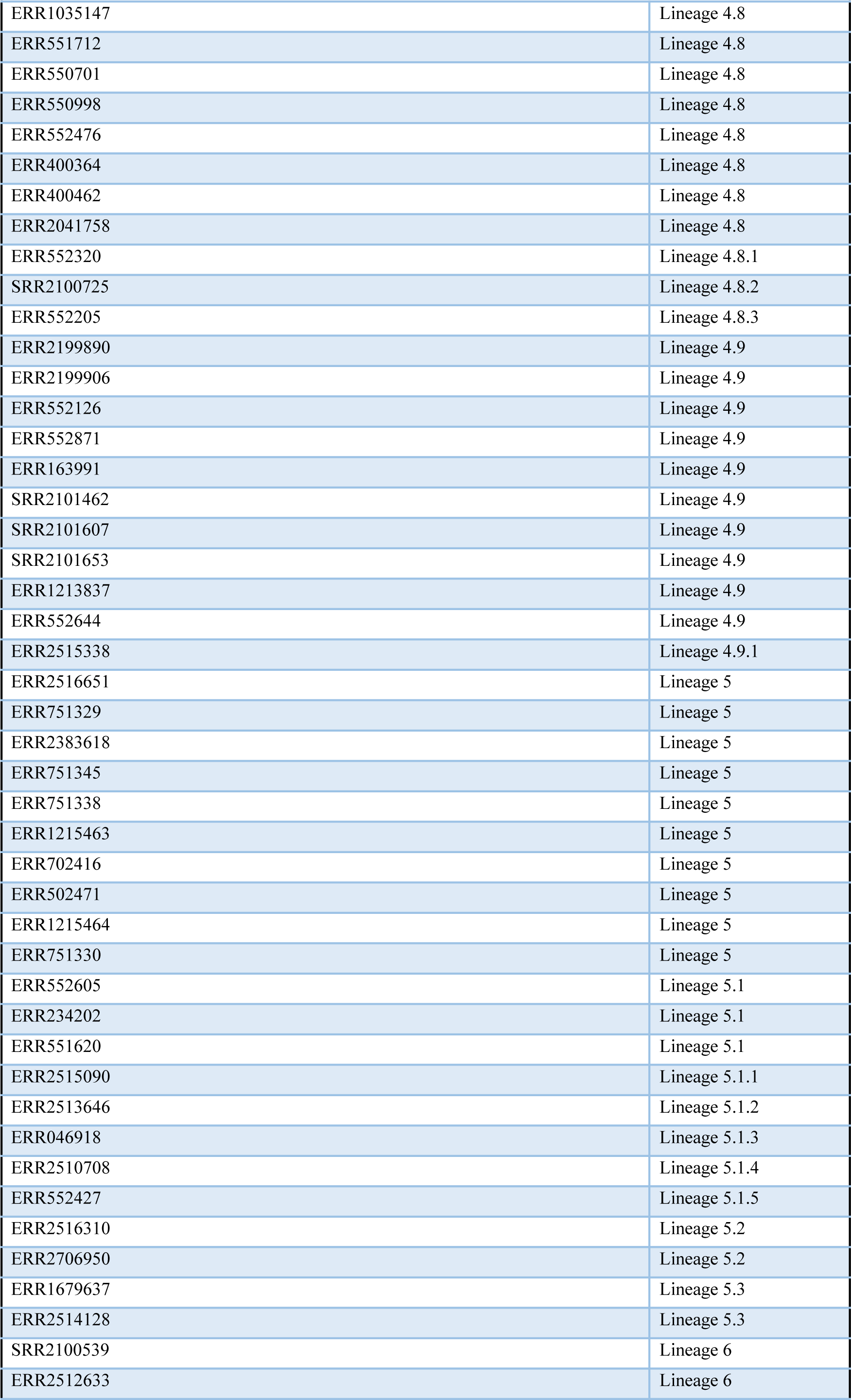
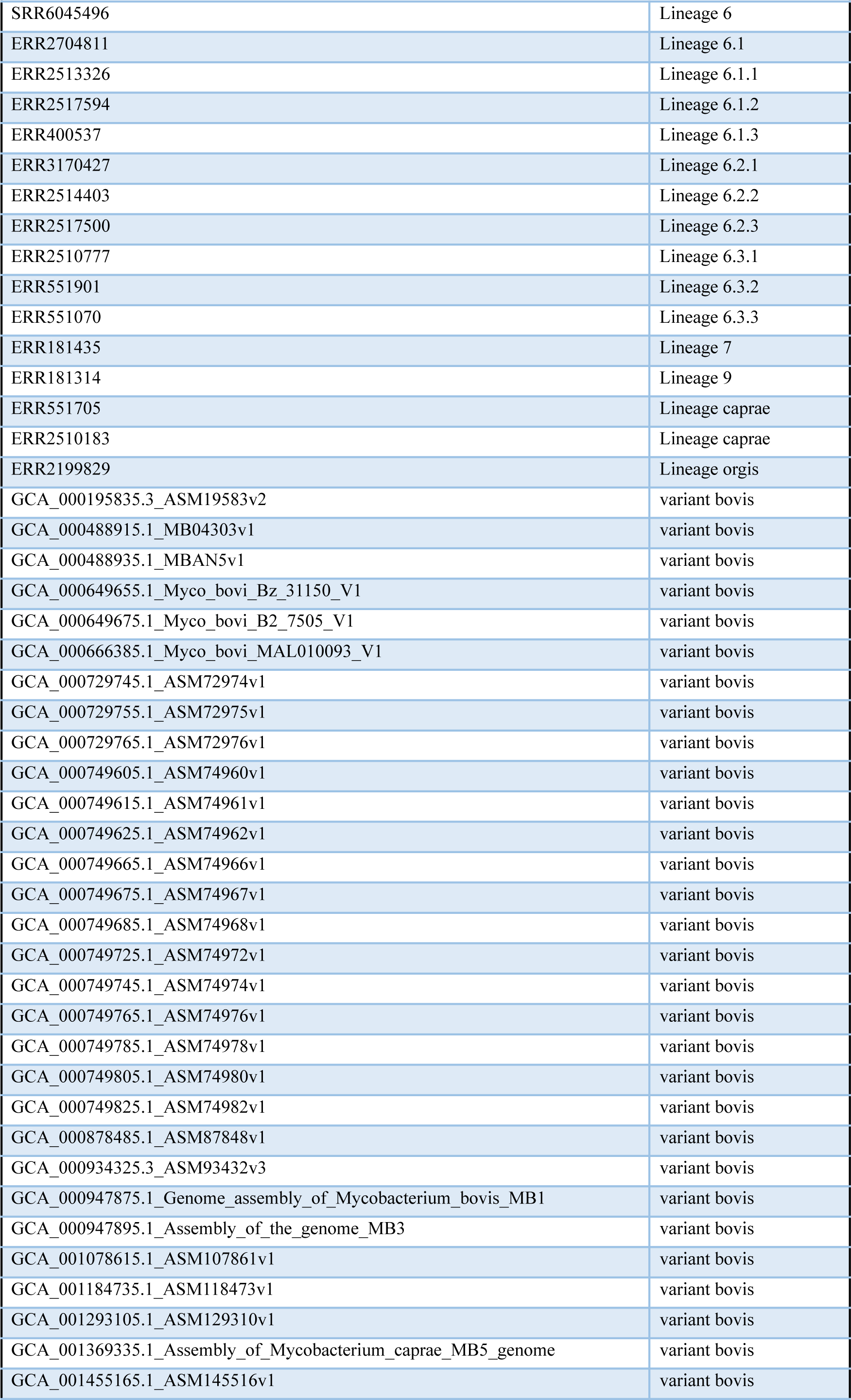
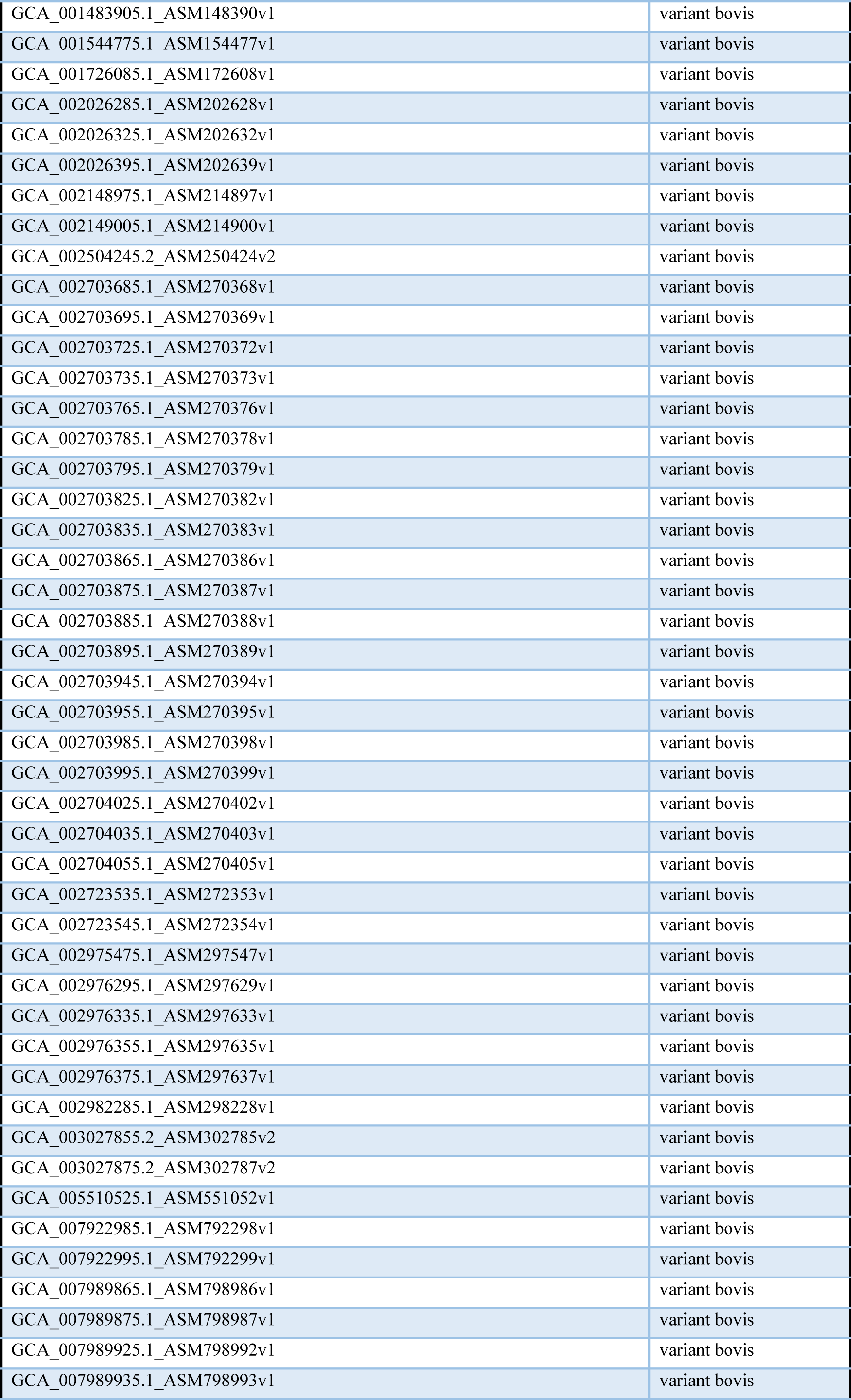
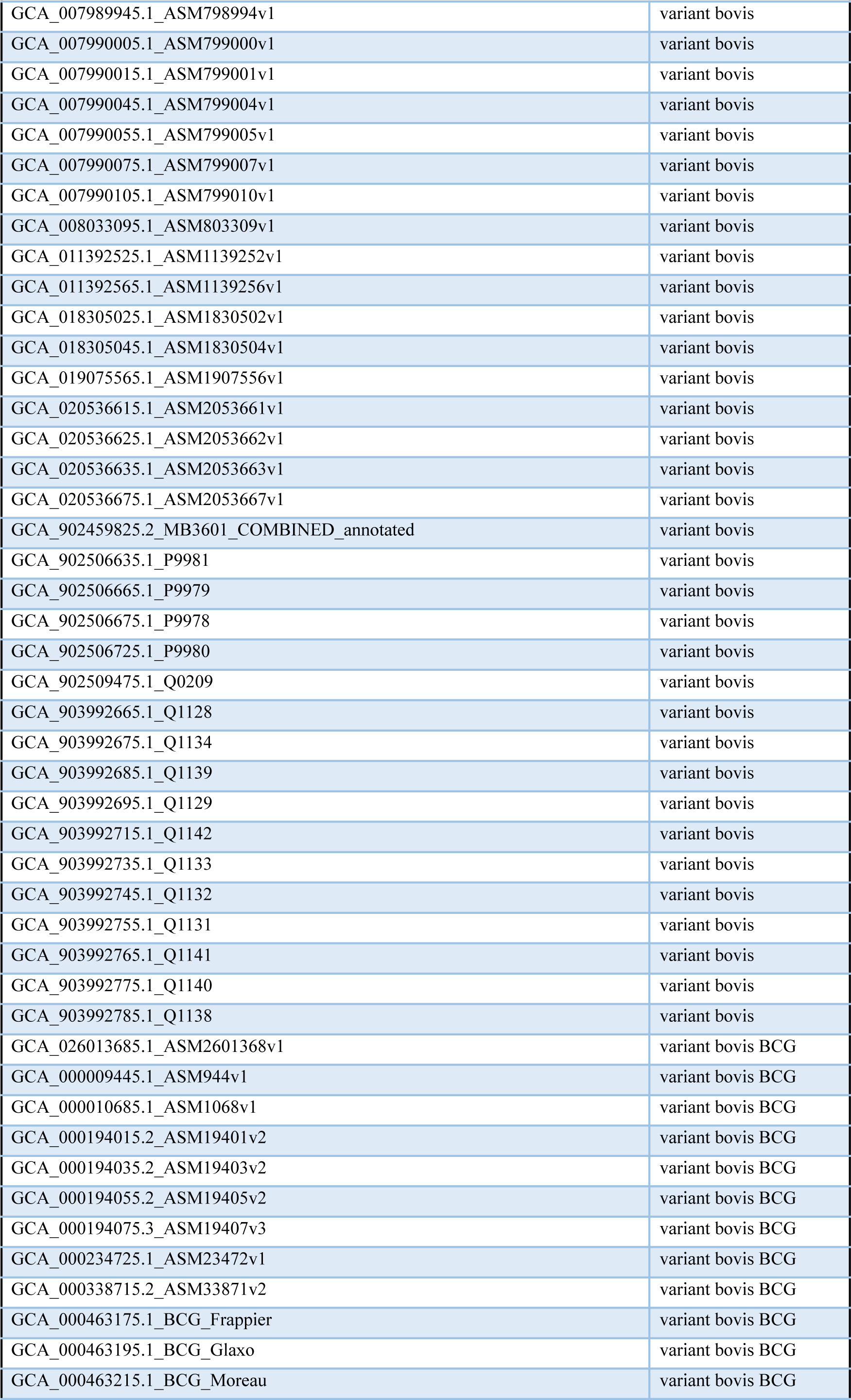
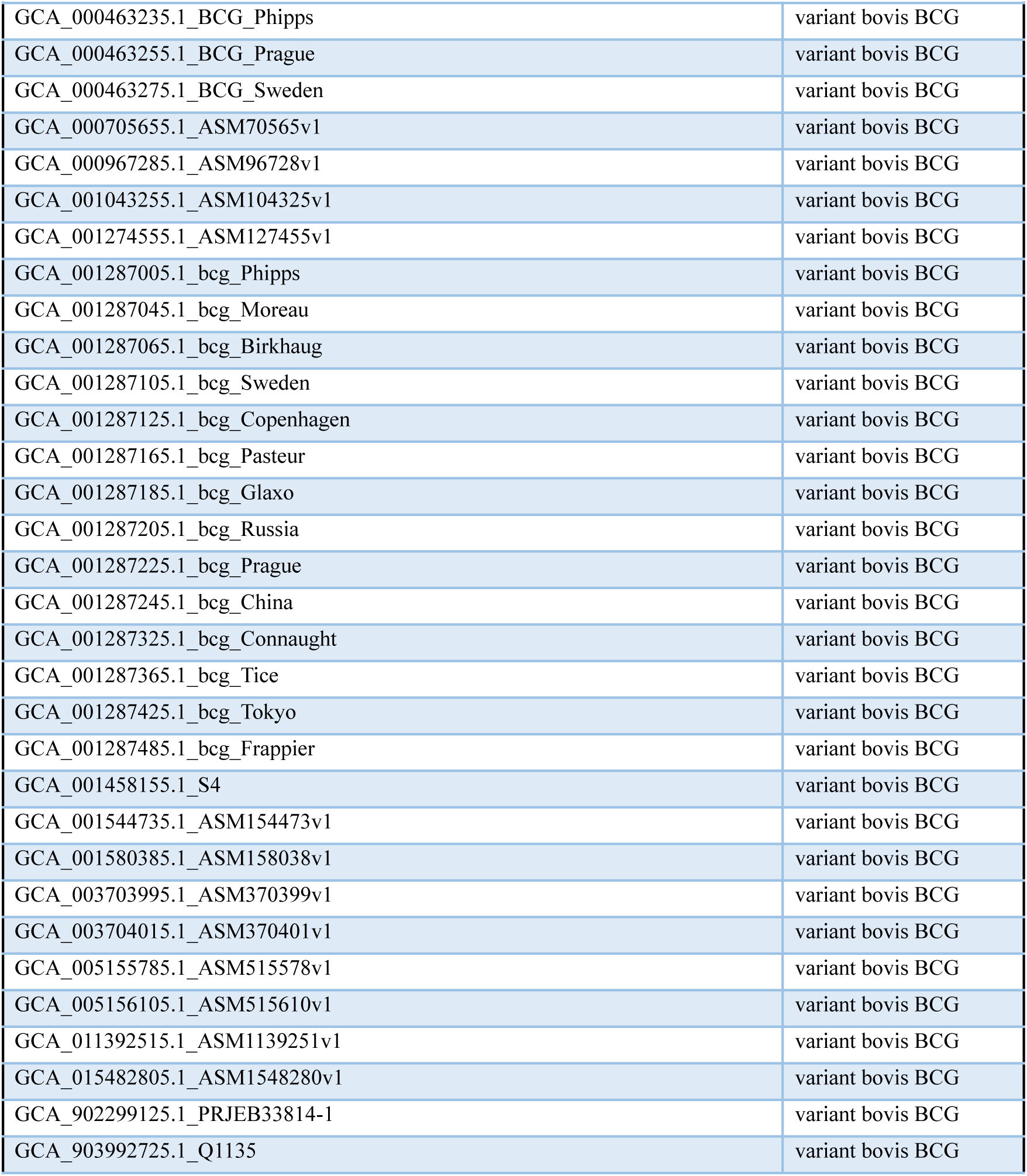
TB lineage dataset.

| Sample ID | Truth Value |
| --- | --- |
| SRR6152881 | Lineage 1 |
| ERR768038 | Lineage 1.1 |
| ERR718556 | Lineage 1.1 |
| ERR718235 | Lineage 1.1 |
| ERR718422 | Lineage 1.1 |
| ERR718288 | Lineage 1.1 |
| ERR718374 | Lineage 1.1 |
| ERR718236 | Lineage 1.1 |
| ERR718262 | Lineage 1.1 |
| ERR718497 | Lineage 1.1 |
| ERR751824 | Lineage 1.1.1 |
| ERR767987 | Lineage 1.1.1.1 |
| ERR2510366 | Lineage 1.1.2 |
| ERR1213947 | Lineage 1.1.2 |
| SRR2100430 | Lineage 1.1.3 |
| SRR2101252 | Lineage 1.1.3.1 |
| ERR2513178 | Lineage 1.1.3.2 |
| ERR2512769 | Lineage 1.1.3.3 |
| ERR2513144 | Lineage 1.1.3.3 |
| ERR553088 | Lineage 1.2 |
| ERR2512840 | Lineage 1.2 |
| ERR400309 | Lineage 1.2 |
| SRR6045034 | Lineage 1.2 |
| ERR2517328 | Lineage 1.2.1 |
| ERR2510788 | Lineage 1.2.2 |
| SRR5067296 | Lineage 1.2.2.1 |
| SRR2100241 | Lineage 1.3.1 |
| ERR2510255 | Lineage 1.3.2 |
| SRR5065560 | Lineage 2.1 |
| ERR552761 | Lineage 2.1 |
| ERR553373 | Lineage 2.1 |
| DRR185119 | Lineage 2.1 |
| DRR184642 | Lineage 2.1 |
| ERR234216 | Lineage 2.1 |
| ERR551044 | Lineage 2.1 |
| ERR181316 | Lineage 2.1 |
| ERR234252 | Lineage 2.1 |
| ERR234248 | Lineage 2.1 |
| ERR502912 | Lineage 2.2 |
| ERR2510746 | Lineage 2.2.1 |
| ERR551871 | Lineage 2.2.1.1 |
| ERR2513866 | Lineage 2.2.1.2 |
| SRR2024907 | Lineage 2.2.2 |
| ERR2514121 | Lineage 3 |
| ERR2514707 | Lineage 3 |
| ERR550978 | Lineage 3 |
| ERR2517625 | Lineage 3 |
| ERR2513473 | Lineage 3 |
| SRR3675535 | Lineage 3 |
| SRR2100827 | Lineage 3 |
| ERR2513685 | Lineage 3 |
| SRR2100044 | Lineage 3 |
| ERR038743 | Lineage 3 |
| ERR2513637 | Lineage 3.1 |
| ERR2514250 | Lineage 3.1 |
| ERR2509882 | Lineage 3.1 |
| ERR551397 | Lineage 3.1 |
| ERR2513554 | Lineage 3.1 |
| SRR2100324 | Lineage 3.1 |
| SRR2100981 | Lineage 3.1 |
| ERR2510590 | Lineage 3.1 |
| ERR2510623 | Lineage 3.1 |
| ERR1034589 | Lineage 3.1 |
| ERR176577 | Lineage 3.1.1 |
| ERR2512807 | Lineage 3.1.2 |
| ERR2512825 | Lineage 3.1.2.1 |
| SRR2100113 | Lineage 3.1.2.2 |
| ERR2200121 | Lineage 3.2 |
| ERR551013 | Lineage 3.2 |
| ERR552820 | Lineage 3.2 |
| SRR6045997 | Lineage 3.2 |
| ERR2514230 | Lineage 3.2 |
| SRR2100025 | Lineage 3.2 |
| SRR2100350 | Lineage 3.2 |
| SRR6045966 | Lineage 3.2 |
| ERR2512416 | Lineage 3.2 |
| ERR2510344 | Lineage 3.2 |
| ERR2199860 | Lineage 4 |
| ERR2517475 | Lineage 4 |
| ERR133897 | Lineage 4 |
| ERR2513995 | Lineage 4 |
| ERR553346 | Lineage 4 |
| ERR2513713 | Lineage 4 |
| ERR552272 | Lineage 4 |
| ERR2516228 | Lineage 4 |
| ERR2514076 | Lineage 4 |
| ERR2517519 | Lineage 4 |
| ERR144613 | Lineage 4.1 |
| ERR2517547 | Lineage 4.1 |
| ERR550633 | Lineage 4.1 |
| SRR2100687 | Lineage 4.1 |
| ERR552879 | Lineage 4.1 |
| SRR2100180 | Lineage 4.1 |
| ERR2517391 | Lineage 4.1 |
| ERR550729 | Lineage 4.1 |
| ERR2513673 | Lineage 4.1.1 |
| SRR2100933 | Lineage 4.1.1.1 |
| ERR2512465 | Lineage 4.1.1.2 |
| ERR2512617 | Lineage 4.1.1.3 |
| SRR2100001 | Lineage 4.1.1.3.1 |
| ERR551893 | Lineage 4.1.2 |
| ERR2512744 | Lineage 4.1.2.1 |
| ERR757170 | Lineage 4.1.2.1.1 |
| ERR552061 | Lineage 4.1.4 |
| ERR751942 | Lineage 4.2 |
| SRR5818596 | Lineage 4.2 |
| SRR6824620 | Lineage 4.2 |
| ERR2179736 | Lineage 4.2 |
| ERR2179737 | Lineage 4.2 |
| ERR2510782 | Lineage 4.2.1 |
| ERR2512872 | Lineage 4.2.1.1 |
| ERR2514985 | Lineage 4.2.2 |
| ERR552483 | Lineage 4.2.2.1 |
| ERR038741 | Lineage 4.2.2.2 |
| ERR551065 | Lineage 4.3 |
| ERR552334 | Lineage 4.3 |
| ERR552704 | Lineage 4.3 |
| ERR551889 | Lineage 4.3 |
| ERR553033 | Lineage 4.3 |
| ERR2510180 | Lineage 4.3 |
| ERR2516611 | Lineage 4.3 |
| ERR2512652 | Lineage 4.3 |
| ERR2517279 | Lineage 4.3 |
| ERR550620 | Lineage 4.3.1 |
| ERR039335 | Lineage 4.3.1.1 |
| SRR2101753 | Lineage 4.3.2 |
| SRR2101516 | Lineage 4.3.2.1 |
| ERR182035 | Lineage 4.3.3 |
| ERR1034794 | Lineage 4.3.4 |
| ERR1035282 | Lineage 4.3.4.1 |
| ERR2514448 | Lineage 4.3.4.2 |
| ERR1034816 | Lineage 4.3.4.2.1 |
| ERR550965 | Lineage 4.4 |
| ERR2514347 | Lineage 4.4 |
| SRR2100805 | Lineage 4.4 |
| ERR2512723 | Lineage 4.4 |
| ERR2514048 | Lineage 4.4 |
| ERR2513635 | Lineage 4.4 |
| ERR1034693 | Lineage 4.4 |
| SRR6045465 | Lineage 4.4 |
| ERR1035311 | Lineage 4.4.1 |
| ERR2516296 | Lineage 4.4.1.1 |
| ERR2512628 | Lineage 4.4.1.1.1 |
| ERR2517511 | Lineage 4.4.1.2 |
| ERR751963 | Lineage 4.4.2 |
| ERR1213839 | Lineage 4.5 |
| ERR1213854 | Lineage 4.5 |
| SRR2100604 | Lineage 4.5 |
| SRR2100720 | Lineage 4.5 |
| ERR1213852 | Lineage 4.5 |
| ERR751714 | Lineage 4.5 |
| SRR671751 | Lineage 4.5 |
| ERR2514763 | Lineage 4.5 |
| SRR671773 | Lineage 4.5 |
| ERR752153 | Lineage 4.5 |
| ERR2512610 | Lineage 4.6 |
| ERR2517343 | Lineage 4.6 |
| SRR2100571 | Lineage 4.6 |
| ERR2510770 | Lineage 4.6 |
| ERR2510807 | Lineage 4.6 |
| ERR551633 | Lineage 4.6 |
| ERR2512531 | Lineage 4.6 |
| SRR2100592 | Lineage 4.6 |
| SRR6046090 | Lineage 4.6 |
| ERR1035142 | Lineage 4.6 |
| ERR181785 | Lineage 4.6.1 |
| ERR553255 | Lineage 4.6.1.1 |
| ERR2513458 | Lineage 4.6.1.2 |
| ERR190355 | Lineage 4.6.2 |
| ERR046838 | Lineage 4.6.2.1 |
| ERR2513583 | Lineage 4.6.2.2 |
| ERR400522 | Lineage 4.6.3 |
| ERR2512590 | Lineage 4.6.4 |
| ERR2514126 | Lineage 4.6.5 |
| SRR2100310 | Lineage 4.7 |
| SRR2100783 | Lineage 4.7 |
| SRR2101046 | Lineage 4.7 |
| SRR2101621 | Lineage 4.7 |
| SRR6045316 | Lineage 4.7 |
| ERR551297 | Lineage 4.7 |
| ERR2517136 | Lineage 4.7 |
| ERR1213922 | Lineage 4.7 |
| ERR2513257 | Lineage 4.7 |
| ERR2516422 | Lineage 4.7 |
| SRR6045004 | Lineage 4.8 |
| ERR1035147 | Lineage 4.8 |
| ERR551712 | Lineage 4.8 |
| ERR550701 | Lineage 4.8 |
| ERR550998 | Lineage 4.8 |
| ERR552476 | Lineage 4.8 |
| ERR400364 | Lineage 4.8 |
| ERR400462 | Lineage 4.8 |
| ERR2041758 | Lineage 4.8 |
| ERR552320 | Lineage 4.8.1 |
| SRR2100725 | Lineage 4.8.2 |
| ERR552205 | Lineage 4.8.3 |
| ERR2199890 | Lineage 4.9 |
| ERR2199906 | Lineage 4.9 |
| ERR552126 | Lineage 4.9 |
| ERR552871 | Lineage 4.9 |
| ERR163991 | Lineage 4.9 |
| SRR2101462 | Lineage 4.9 |
| SRR2101607 | Lineage 4.9 |
| SRR2101653 | Lineage 4.9 |
| ERR1213837 | Lineage 4.9 |
| ERR552644 | Lineage 4.9 |
| ERR2515338 | Lineage 4.9.1 |
| ERR2516651 | Lineage 5 |
| ERR751329 | Lineage 5 |
| ERR2383618 | Lineage 5 |
| ERR751345 | Lineage 5 |
| ERR751338 | Lineage 5 |
| ERR1215463 | Lineage 5 |
| ERR702416 | Lineage 5 |
| ERR502471 | Lineage 5 |
| ERR1215464 | Lineage 5 |
| ERR751330 | Lineage 5 |
| ERR552605 | Lineage 5.1 |
| ERR234202 | Lineage 5.1 |
| ERR551620 | Lineage 5.1 |
| ERR2515090 | Lineage 5.1.1 |
| ERR2513646 | Lineage 5.1.2 |
| ERR046918 | Lineage 5.1.3 |
| ERR2510708 | Lineage 5.1.4 |
| ERR552427 | Lineage 5.1.5 |
| ERR2516310 | Lineage 5.2 |
| ERR2706950 | Lineage 5.2 |
| ERR1679637 | Lineage 5.3 |
| ERR2514128 | Lineage 5.3 |
| SRR2100539 | Lineage 6 |
| ERR2512633 | Lineage 6 |
| SRR6045496 | Lineage 6 |
| ERR2704811 | Lineage 6.1 |
| ERR2513326 | Lineage 6.1.1 |
| ERR2517594 | Lineage 6.1.2 |
| ERR400537 | Lineage 6.1.3 |
| ERR3170427 | Lineage 6.2.1 |
| ERR2514403 | Lineage 6.2.2 |
| ERR2517500 | Lineage 6.2.3 |
| ERR2510777 | Lineage 6.3.1 |
| ERR551901 | Lineage 6.3.2 |
| ERR551070 | Lineage 6.3.3 |
| ERR181435 | Lineage 7 |
| ERR181314 | Lineage 9 |
| ERR551705 | Lineage caprae |
| ERR2510183 | Lineage caprae |
| ERR2199829 | Lineage orgis |
| GCA_000195835.3_ASM19583v2 | variant bovis |
| GCA_000488915.1_MB04303v1 | variant bovis |
| GCA_000488935.1_MBAN5v1 | variant bovis |
| GCA_000649655.1_Myco_bovi_Bz_31150_V1 | variant bovis |
| GCA_000649675.1_Myco_bovi_B2_7505_V1 | variant bovis |
| GCA_000666385.1_Myco_bovi_MAL010093_V1 | variant bovis |
| GCA_000729745.1_ASM72974v1 | variant bovis |
| GCA_000729755.1_ASM72975v1 | variant bovis |
| GCA_000729765.1_ASM72976v1 | variant bovis |
| GCA_000749605.1_ASM74960v1 | variant bovis |
| GCA_000749615.1_ASM74961v1 | variant bovis |
| GCA_000749625.1_ASM74962v1 | variant bovis |
| GCA_000749665.1_ASM74966v1 | variant bovis |
| GCA_000749675.1_ASM74967v1 | variant bovis |
| GCA_000749685.1_ASM74968v1 | variant bovis |
| GCA_000749725.1_ASM74972v1 | variant bovis |
| GCA_000749745.1_ASM74974v1 | variant bovis |
| GCA_000749765.1_ASM74976v1 | variant bovis |
| GCA_000749785.1_ASM74978v1 | variant bovis |
| GCA_000749805.1_ASM74980v1 | variant bovis |
| GCA_000749825.1_ASM74982v1 | variant bovis |
| GCA_000878485.1_ASM87848v1 | variant bovis |
| GCA_000934325.3_ASM93432v3 | variant bovis |
| GCA_000947875.1_Genome_assembly_of_Mycobacterium_bovis_MB1 | variant bovis |
| GCA_000947895.1_Assembly_of_the_genome_MB3 | variant bovis |
| GCA_001078615.1_ASM107861v1 | variant bovis |
| GCA_001184735.1_ASM118473v1 | variant bovis |
| GCA_001293105.1_ASM129310v1 | variant bovis |
| GCA_001369335.1_Assembly_of_Mycobacterium_caprae_MB5_genome | variant bovis |
| GCA_001455165.1_ASM145516v1 | variant bovis |
| GCA_001483905.1_ASM148390v1 | variant bovis |
| GCA_001544775.1_ASM154477v1 | variant bovis |
| GCA_001726085.1_ASM172608v1 | variant bovis |
| GCA_002026285.1_ASM202628v1 | variant bovis |
| GCA_002026325.1_ASM202632v1 | variant bovis |
| GCA_002026395.1_ASM202639v1 | variant bovis |
| GCA_002148975.1_ASM214897v1 | variant bovis |
| GCA_002149005.1_ASM214900v1 | variant bovis |
| GCA_002504245.2_ASM250424v2 | variant bovis |
| GCA_002703685.1_ASM270368v1 | variant bovis |
| GCA_002703695.1_ASM270369v1 | variant bovis |
| GCA_002703725.1_ASM270372v1 | variant bovis |
| GCA_002703735.1_ASM270373v1 | variant bovis |
| GCA_002703765.1_ASM270376v1 | variant bovis |
| GCA_002703785.1_ASM270378v1 | variant bovis |
| GCA_002703795.1_ASM270379v1 | variant bovis |
| GCA_002703825.1_ASM270382v1 | variant bovis |
| GCA_002703835.1_ASM270383v1 | variant bovis |
| GCA_002703865.1_ASM270386v1 | variant bovis |
| GCA_002703875.1_ASM270387v1 | variant bovis |
| GCA_002703885.1_ASM270388v1 | variant bovis |
| GCA_002703895.1_ASM270389v1 | variant bovis |
| GCA_002703945.1_ASM270394v1 | variant bovis |
| GCA_002703955.1_ASM270395v1 | variant bovis |
| GCA_002703985.1_ASM270398v1 | variant bovis |
| GCA_002703995.1_ASM270399v1 | variant bovis |
| GCA_002704025.1_ASM270402v1 | variant bovis |
| GCA_002704035.1_ASM270403v1 | variant bovis |
| GCA_002704055.1_ASM270405v1 | variant bovis |
| GCA_002723535.1_ASM272353v1 | variant bovis |
| GCA_002723545.1_ASM272354v1 | variant bovis |
| GCA_002975475.1_ASM297547v1 | variant bovis |
| GCA_002976295.1_ASM297629v1 | variant bovis |
| GCA_002976335.1_ASM297633v1 | variant bovis |
| GCA_002976355.1_ASM297635v1 | variant bovis |
| GCA_002976375.1_ASM297637v1 | variant bovis |
| GCA_002982285.1_ASM298228v1 | variant bovis |
| GCA_003027855.2_ASM302785v2 | variant bovis |
| GCA_003027875.2_ASM302787v2 | variant bovis |
| GCA_005510525.1_ASM551052v1 | variant bovis |
| GCA_007922985.1_ASM792298v1 | variant bovis |
| GCA_007922995.1_ASM792299v1 | variant bovis |
| GCA_007989865.1_ASM798986v1 | variant bovis |
| GCA_007989875.1_ASM798987v1 | variant bovis |
| GCA_007989925.1_ASM798992v1 | variant bovis |
| GCA_007989935.1_ASM798993v1 | variant bovis |
| GCA_007989945.1_ASM798994v1 | variant bovis |
| GCA_007990005.1_ASM799000v1 | variant bovis |
| GCA_007990015.1_ASM799001v1 | variant bovis |
| GCA_007990045.1_ASM799004v1 | variant bovis |
| GCA_007990055.1_ASM799005v1 | variant bovis |
| GCA_007990075.1_ASM799007v1 | variant bovis |
| GCA_007990105.1_ASM799010v1 | variant bovis |
| GCA_008033095.1_ASM803309v1 | variant bovis |
| GCA_011392525.1_ASM1139252v1 | variant bovis |
| GCA_011392565.1_ASM1139256v1 | variant bovis |
| GCA_018305025.1_ASM1830502v1 | variant bovis |
| GCA_018305045.1_ASM1830504v1 | variant bovis |
| GCA_019075565.1_ASM1907556v1 | variant bovis |
| GCA_020536615.1_ASM2053661v1 | variant bovis |
| GCA_020536625.1_ASM2053662v1 | variant bovis |
| GCA_020536635.1_ASM2053663v1 | variant bovis |
| GCA_020536675.1_ASM2053667v1 | variant bovis |
| GCA_902459825.2_MB3601_COMBINED_annotated | variant bovis |
| GCA_902506635.1_P9981 | variant bovis |
| GCA_902506665.1_P9979 | variant bovis |
| GCA_902506675.1_P9978 | variant bovis |
| GCA_902506725.1_P9980 | variant bovis |
| GCA_902509475.1_Q0209 | variant bovis |
| GCA_903992665.1_Q1128 | variant bovis |
| GCA_903992675.1_Q1134 | variant bovis |
| GCA_903992685.1_Q1139 | variant bovis |
| GCA_903992695.1_Q1129 | variant bovis |
| GCA_903992715.1_Q1142 | variant bovis |
| GCA_903992735.1_Q1133 | variant bovis |
| GCA_903992745.1_Q1132 | variant bovis |
| GCA_903992755.1_Q1131 | variant bovis |
| GCA_903992765.1_Q1141 | variant bovis |
| GCA_903992775.1_Q1140 | variant bovis |
| GCA_903992785.1_Q1138 | variant bovis |
| GCA_026013685.1_ASM2601368v1 | variant bovis BCG |
| GCA_000009445.1_ASM944v1 | variant bovis BCG |
| GCA_000010685.1_ASM1068v1 | variant bovis BCG |
| GCA_000194015.2_ASM19401v2 | variant bovis BCG |
| GCA_000194035.2_ASM19403v2 | variant bovis BCG |
| GCA_000194055.2_ASM19405v2 | variant bovis BCG |
| GCA_000194075.3_ASM19407v3 | variant bovis BCG |
| GCA_000234725.1_ASM23472v1 | variant bovis BCG |
| GCA_000338715.2_ASM33871v2 | variant bovis BCG |
| GCA_000463175.1_BCG_Frappier | variant bovis BCG |
| GCA_000463195.1_BCG_Glaxo | variant bovis BCG |
| GCA_000463215.1_BCG_Moreau | variant bovis BCG |
| GCA_000463235.1_BCG_Phipps | variant bovis BCG |
| GCA_000463255.1_BCG_Prague | variant bovis BCG |
| GCA_000463275.1_BCG_Sweden | variant bovis BCG |
| GCA_000705655.1_ASM70565v1 | variant bovis BCG |
| GCA_000967285.1_ASM96728v1 | variant bovis BCG |
| GCA_001043255.1_ASM104325v1 | variant bovis BCG |
| GCA_001274555.1_ASM127455v1 | variant bovis BCG |
| GCA_001287005.1_bcg_Phipps | variant bovis BCG |
| GCA_001287045.1_bcg_Moreau | variant bovis BCG |
| GCA_001287065.1_bcg_Birkhaug | variant bovis BCG |
| GCA_001287105.1_bcg_Sweden | variant bovis BCG |
| GCA_001287125.1_bcg_Copenhagen | variant bovis BCG |
| GCA_001287165.1_bcg_Pasteur | variant bovis BCG |
| GCA_001287185.1_bcg_Glaxo | variant bovis BCG |
| GCA_001287205.1_bcg_Russia | variant bovis BCG |
| GCA_001287225.1_bcg_Prague | variant bovis BCG |
| GCA_001287245.1_bcg_China | variant bovis BCG |
| GCA_001287325.1_bcg_Connaught | variant bovis BCG |
| GCA_001287365.1_bcg_Tice | variant bovis BCG |
| GCA_001287425.1_bcg_Tokyo | variant bovis BCG |
| GCA_001287485.1_bcg_Frappier | variant bovis BCG |
| GCA_001458155.1_S4 | variant bovis BCG |
| GCA_001544735.1_ASM154473v1 | variant bovis BCG |
| GCA_001580385.1_ASM158038v1 | variant bovis BCG |
| GCA_003703995.1_ASM370399v1 | variant bovis BCG |
| GCA_003704015.1_ASM370401v1 | variant bovis BCG |
| GCA_005155785.1_ASM515578v1 | variant bovis BCG |
| GCA_005156105.1_ASM515610v1 | variant bovis BCG |
| GCA_011392515.1_ASM1139251v1 | variant bovis BCG |
| GCA_015482805.1_ASM1548280v1 | variant bovis BCG |
| GCA_902299125.1_PRJEB33814-1 | variant bovis BCG |
| GCA_903992725.1_Q1135 | variant bovis BCG |

**Table S4.** Result frequency of running the lineage dataset through each profiler.

|  |  | PASS | Overqualified | Underqualified | Type 1 FAIL | Type 2 FAIL | Other |
| --- | --- | --- | --- | --- | --- | --- | --- |
| *L1*<br>(*n*=28) | Afanc | 0.643 | 0.179 | 0.000 | 0.143 | 0.000 | 0.036 |
|  | Tbprofiler | 0.643 | 0.179 | 0.000 | 0.143 | 0.000 | 0.036 |
|  | Mykrobe | 0.571 | 0.143 | 0.107 | 0.143 | 0.036 | 0.000 |
| *L2*<br>(*n*=15) | Afanc | 0.933 | 0.067 | 0.000 | 0.000 | 0.000 | 0.000 |
|  | Tbprofiler | 0.933 | 0.067 | 0.000 | 0.000 | 0.000 | 0.000 |
|  | Mykrobe | 0.733 | 0.000 | 0.067 | 0.200 | 0.000 | 0.000 |
| *L3*<br>(*n=34*) | Afanc | 1.000 | 0.000 | 0.000 | 0.000 | 0.000 | 0.000 |
|  | Tbprofiler | 0.706 | 0.000 | 0.000 | 0.294 | 0.000 | 0.000 |
|  | Mykrobe | 0.412 | 0.000 | 0.588 | 0.000 | 0.000 | 0.000 |
| *L4*<br>(*n=130*) | Afanc | 0.977 | 0.008 | 0.008 | 0.000 | 0.000 | 0.008 |
|  | Tbprofiler | 0.985 | 0.000 | 0.000 | 0.000 | 0.008 | 0.008 |
|  | Mykrobe | 0.592 | 0.085 | 0.046 | 0.277 | 0.000 | 0.000 |
| *L5*<br>(*n=22*) | Afanc | 0.818 | 0.045 | 0.136 | 0.000 | 0.000 | 0.000 |
|  | Tbprofiler | 0.727 | 0.273 | 0.000 | 0.000 | 0.000 | 0.000 |
|  | Mykrobe | 0.455 | 0.000 | 0.545 | 0.000 | 0.000 | 0.000 |
| *L6*<br>(*n=12*) | Afanc | 0.917 | 0.083 | 0.000 | 0.000 | 0.000 | 0.000 |
|  | Tbprofiler | 0.833 | 0.083 | 0.083 | 0.000 | 0.000 | 0.000 |
|  | Mykrobe | 0.250 | 0.000 | 0.750 | 0.000 | 0.000 | 0.000 |
| *L7*<br>(*n=1*) | Afanc | 1.000 | 0.000 | 0.000 | 0.000 | 0.000 | 0.000 |
|  | Tbprofiler | 1.000 | 0.000 | 0.000 | 0.000 | 0.000 | 0.000 |
|  | Mykrobe | 1.000 | 0.000 | 0.000 | 0.000 | 0.000 | 0.000 |
| *L9*<br>(*n=1*) | Afanc | 1.000 | 0.000 | 0.000 | 0.000 | 0.000 | 0.000 |
|  | Tbprofiler | 1.000 | 0.000 | 0.000 | 0.000 | 0.000 | 0.000 |
|  | Mykrobe | 0.000 | 0.000 | 0.000 | 0.000 | 1.000 | 0.000 |
| *caprae*<br>(*n=2*) | Afanc | 1.000 | 0.000 | 0.000 | 0.000 | 0.000 | 0.000 |
|  | Tbprofiler | 1.000 | 0.000 | 0.000 | 0.000 | 0.000 | 0.000 |
|  | Mykrobe | 1.000 | 0.000 | 0.000 | 0.000 | 0.000 | 0.000 |
| *orgis*<br>(*n=1*) | Afanc | 1.000 | 0.000 | 0.000 | 0.000 | 0.000 | 0.000 |
|  | Tbprofiler | 1.000 | 0.000 | 0.000 | 0.000 | 0.000 | 0.000 |
|  | Mykrobe | 0.000 | 0.000 | 0.000 | 0.000 | 1.000 | 0.000 |
| *bovis*<br>(*n=110*) | Afanc | 0.909 | 0.000 | 0.000 | 0.073 | 0.000 | 0.018 |
|  | Tbprofiler | 0.909 | 0.000 | 0.000 | 0.064 | 0.027 | 0.000 |
|  | Mykrobe | 0.918 | 0.000 | 0.000 | 0.055 | 0.009 | 0.018 |
| *bovis*<br>*BCG*<br>(*n*=44) | Afanc | 0.977 | 0.000 | 0.000 | 0.000 | 0.023 | 0.000 |
|  | Tbprofiler | 0.977 | 0.000 | 0.000 | 0.000 | 0.023 | 0.000 |
|  | Mykrobe | 0.955 | 0.000 | 0.000 | 0.000 | 0.045 | 0.000 |
| *Total*<br>(*n*=400) | Afanc | 0.925 | 0.023 | 0.010 | 0.030 | 0.003 | 0.010 |
|  | Tbprofiler | 0.895 | 0.033 | 0.003 | 0.053 | 0.013 | 0.005 |
|  | Mykrobe | 0.693 | 0.038 | 0.128 | 0.123 | 0.015 | 0.005 |

## Bibliography

Breitwieser, F.P., Baker, D.N., Salzberg, S.L., 2018. KrakenUniq: confident and fast metagenomics classification using unique k-mer counts. Genome Biology 19. 10.1186/s13059-018-1568-0

Gupta, R.S., Lo, B., Son, J., 2018. Phylogenomics and Comparative Genomic Studies Robustly Support Division of the Genus Mycobacterium into an Emended Genus Mycobacterium and Four Novel Genera. Frontiers in Microbiology 9. 10.3389/fmicb.2018.00067

Houben, R.M.G.J., Dodd, P.J., 2016. The Global Burden of Latent Tuberculosis Infection: A Re-estimation Using Mathematical Modelling. PLOS Medicine 13, e1002152. 10.1371/journal.pmed.1002152

Huang, W., Li, L., Myers, J.R., Marth, G.T., 2011. ART: a next-generation sequencing read simulator. Bioinformatics 28, 593–594. 10.1093/bioinformatics/btr708

Hunt, M., Bradley, P., Lapierre, S.G., Heys, S., Thomsit, M., Hall, M.B., Malone, K.M., Wintringer, P., Walker, T.M., Cirillo, D.M., Comas, I., Farhat, M.R., Fowler, P., Gardy, J., Ismail, N., Kohl, T.A., Mathys, V., Merker, M., Niemann, S., Omar, S.V., Sintchenko, V., Smith, G., Soolingen, D. van, Supply, P., Tahseen, S., Wilcox, M., Arandjelovic, I., Peto, T.E.A., Crook, D.W., Iqbal, Z., 2019. Antibiotic resistance prediction for Mycobacterium tuberculosis from genome sequence data with Mykrobe. Wellcome Open Research 4, 191. 10.12688/wellcomeopenres.15603.1

Lu, J., Breitwieser, F.P., Thielen, P., Salzberg, S.L., 2017. Bracken: estimating species abundance in metagenomics data. PeerJ Computer Science 3, e104. 10.7717/peerj-cs.104

Napier, G., Campino, S., Merid, Y., Abebe, M., Woldeamanuel, Y., Aseffa, A., Hibberd, M.L., Phelan, J., Clark, T.G., 2020. Robust barcoding and identification of Mycobacterium tuberculosis lineages for epidemiological and clinical studies. Genome Medicine 12. 10.1186/s13073-020-00817-3

Ondov, B.D., Treangen, T.J., Melsted, P., Mallonee, A.B., Bergman, N.H., Koren, S., Phillippy, A.M., 2016. Mash: fast genome and metagenome distance estimation using MinHash. Genome Biology 17. 10.1186/s13059-016-0997-x

Ratnatunga, C.N., Lutzky, V.P., Kupz, A., Doolan, D.L., Reid, D.W., Field, M., Bell, S.C., Thomson, R.M., Miles, J.J., 2020. The Rise of Non-Tuberculosis Mycobacterial Lung Disease. Frontiers in Immunology 11. 10.3389/fimmu.2020.00303

Wood, D.E., Lu, J., Langmead, B., 2019. Improved metagenomic analysis with Kraken 2. Genome Biology 20. 10.1186/s13059-019-1891-0

Wood, D.E., Salzberg, S.L., 2014. Kraken: ultrafast metagenomic sequence classification using exact alignments. Genome Biology 15. 10.1186/gb-2014-15-3-r46

